# Signatures of Mutational Processes in Human DNA Evolution

**DOI:** 10.1101/2021.01.09.426041

**Authors:** Hamid Hamidi, Hamid Alinejad-Rokny, Tim Coorens, Rashesh Sanghvi, Sarah J Lindsay, Raheleh Rahbari, Diako Ebrahimi

## Abstract

The human genome contains over 100 million SNPs, most of which are C/T (G/A) variations. The type and sequence context of these SNPs are not random, suggesting that they are caused by distinct mutational processes. Deciphering the mutational signatures is a crucial step to discovering the molecular processes responsible for DNA variations across human populations, and potentially for causing genetic diseases. Our analyses of the 1000 Genomes Project SNPs and germline *de novo* mutations suggest that at least four mutational processes are responsible for human genetic variations. One process is European-specific and no longer active. The remaining three processes are currently active in all human populations. Two of the active processes co-occur and leave a single joint mutational signature in human nuclear DNA. The third active process is specific to mitochondrial DNA, and inflicts C-to-T mutations at mostly non-CG sites. We found neither evidence of APOBEC-induced cytosine deamination in the human germline, nor *de novo* mutation enrichment within certain regions of the human genome.

## Introduction

Variations observed in the human genome are signatures of contemporary and historic mutational processes. A systematic analysis of these genetic variations can help shed light on the underlying mutational processes and their potential roles as risk factors for diseases^1^. There is currently no consensus on the type and proportion of mutational signatures in human populations. Comparison of the 1000 Genomes Project^2^ SNPs in different populations revealed an elevated rate of TCC>TTC variations among Europeans^3,4^. However, analysis of the same dataset using hierarchical clustering showed three distinct patterns of mutation enrichment among populations^5^. Moreover, a recent study in which genomic coordinates of rare mutations were taken into consideration revealed 14 unique patterns of mutations in the TOPMed dataset^6^. Unexpectedly, few prior studies report signatures of germline mutations induced by proteins of our innate immune system known as APOBEC (Apolipoprotein B mRNA editing enzyme, catalytic polypeptide)^7–9^. Members of this enzyme family such as APOBEC3A and APOBEC3B are considered to be the primary sources of cancer mutational signatures known as SBS2 and SBS13^10–12^. A study of rare SNPs in the 1000 Genomes Project reported that APOBEC3A and APOBEC3B were likely responsible for 20% of the germline C>T and C>G mutations within TCW (W:T, A)^8^. Pinto *et al*. reached a similar conclusion but suggested that a cytoplasmic and antiviral member of the APOBEC3 family, APOBEC3G, was the source of germline mutations and primate evolution^7^. Other studies reported clusters of C>G mutations but ruled out a role for APOBEC3 because the context of mutations was not enriched for TCW^13,14,15^. The methodologies used in these studies often assume that each mutation type (e.g. TCC>TTC) is generated by only one mutational process. In other words, different mutational processes cannot generate the same mutation. Studies of cancer mutations suggest that this assumption is often incorrect. For example, CCC>CAC can have several different sources, such as defective homologous recombination-based DNA damage repair (SBS3), tobacco smoking (SBS4), or even exposure to aflatoxin (SBS24)^11^. An alternative approach used to identify mutational signatures without making this assumption is based on the deconvolution of mutational datasets^10,11,16,17^. Recent quantitative analysis approaches, based on non-negative matrix factorization (NMF)^18^ of complex tumor mutations, have been able to decipher signatures of many mutational processes in cancer^10,16^. In these studies, each mutational signature is defined by a measure of the relative frequency of 96 mutation types (nNn>nMn, where N is mutated to M and n: A, C, G, or T). These studies have led to invaluable findings about the mechanisms underlying mutations in cancer such as spontaneous deamination of 5-methylcytosine (SBS1), APOBEC mutagenesis (SBS2 and SBS13), and homologous recombination-based repair deficiency due to BRCA mutations (SBS3)^10,11,19^. NMF and similar mutation deconvolution approaches have also been used recently to gain insight into mutational processes driving human DNA evolution^6,17^. For example, Mathieson *et al*. used independent component analysis followed by NMF to deconvolute signatures of rare mutations. By analyzing two independent datasets, high coverage sequences of 300 individuals from the Simons Genome Diversity Project and phase 3 of the 1000 Genomes Project, the authors discovered four mutation signatures^17^.

A study of multi-sibling families revealed that the spectrum of *de novo* germline mutations can be explained by only two signatures, SBS1 and SBS5, which are found ubiquitously in cancer^20^. SBS1 is referred to as ‘aging’ signature and is thought to be a signature of CG (a.k.a. CpG) methylations followed by spontaneous deamination^10,21^. This signature is characterized by four C>T mutation peaks at CG sites. Signature SBS5, which is also associated with aging, has an unknown etiology and is characterized by C>T and T>C mutations, with the latter showing enrichment for ATA, ATG, and ATT motifs.

In this study, we performed a systematic analysis of population and trios datasets using four independent approaches, namely, PCA (principal component analysis), NPCA (nonnegative PCA)^22^, NMF (non-negative matrix factorization), and HDP (hierarchical Dirichlet process)^23–26^ to identify signatures of present and past mutational processes in different human populations. Our analyses revealed that present-day *de novo* germline mutations can be described using a single mutational signature, which is population independent. We also identified a European-specific signature reported previously^3–5,17,27,28^ and a novel mutational signature that is specific to mitochondria DNA.

## Results

### Deciphering mutational signatures

#### 1000 Genomes Project Dataset

We quantified the abundance of all 96 types of mutations (nNn>nMn) in the 1000 Genomes Project donors by comparing them to the most common human ancestral sequence and choosing the low frequency SNPs (<0.01) (Method). PCA applied to this raw mutation data matrix (2,504 × 96) showed four PCs (**Supplementary Fig. S1**). **Fig. 1** shows the distribution of populations in PC1-PC2 and PC3-PC4 spaces.

**Fig 1.**
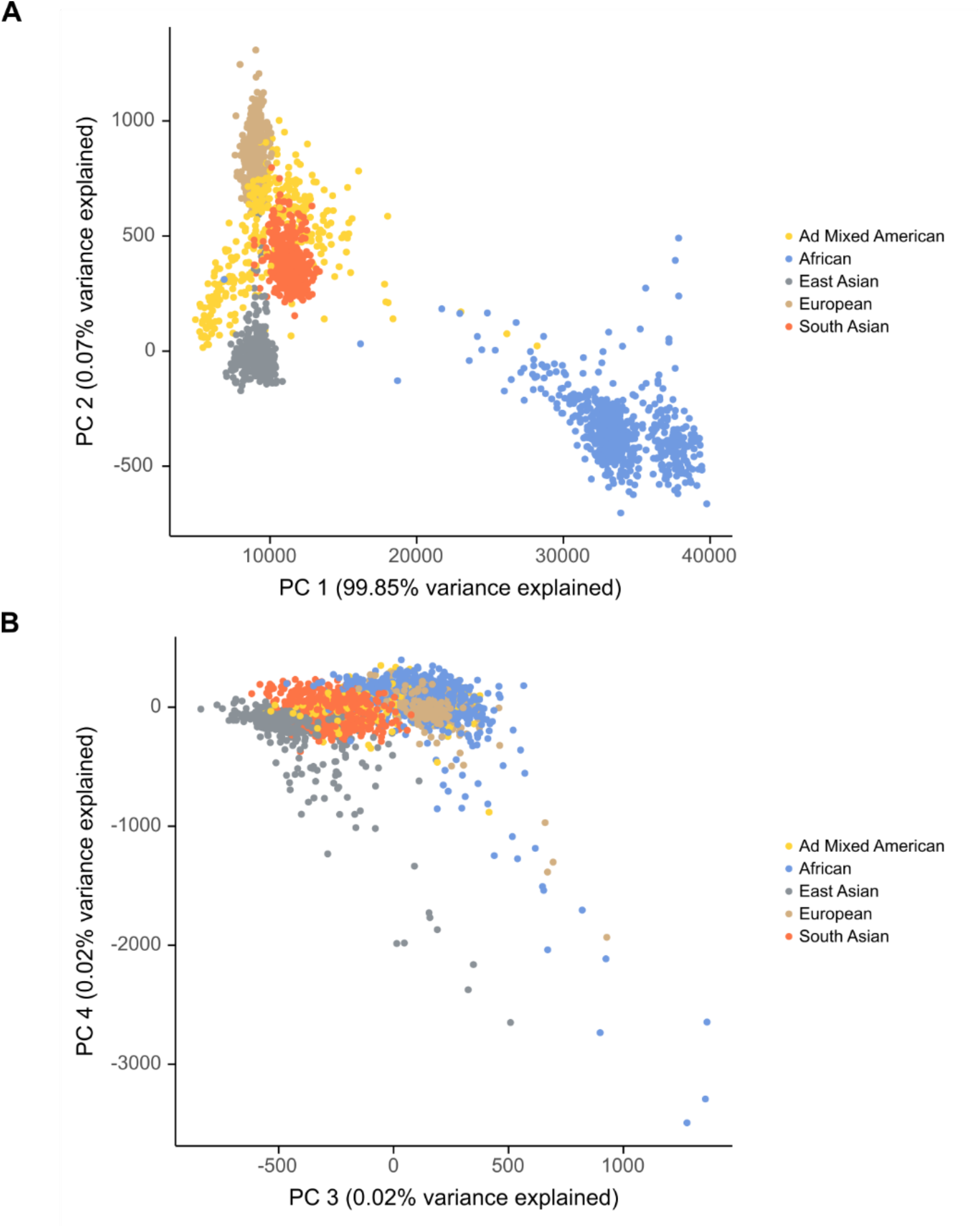
PCA applied to raw mutation data matrix. (A) PC1-PC2 (B) PC3-PC4

Analysis of rare SNPs using NPCA revealed four significant principal components (i.e. mutational signatures), explaining ∼70% of total variance (**Supplementary Fig. S2**). The first principal component is characterized by, mostly, C>T and T>C mutations (**Fig. 2A**). This signature (1000G-Sig1) has four main C>T peaks at CG motifs (ACG, CCG, GCG, and TCG) and one C>T peak at ACA. Additionally, it has three main T>C peaks at ATA, ATG, and ATT. The second mutational signature (1000G-Sig2) is represented by multiple C>T peaks, the top three being at motifs A<u>C</u>C, T<u>C</u>C, and T<u>C</u>T (**Fig. 2A**). The third mutational signature (1000G-Sig3) has five main peaks of C>A at ACA, C>T at TCT, T>A at ATA, and T>G at GTG and GTT (**Fig. 2A**). The fourth mutation signature (1000G-Sig4) is represented by multiple C>G and C>T peaks mostly at non-CG sites and four main T>C peaks at ATA, ATG, ATT, and TTT (**Fig. 2A**). Analysis of the same mutation matrix using NMF also returned four mutational signatures (**Supplementary Fig. S3**). Each NMF mutational signature shows its unique set of peaks, which are identical to those of NPCA signatures after a background correction (Method). The cosine similarities between the mutational signatures obtained using NMF and their corresponding signature in NPCA were >0.82 (**Supplementary Fig. S4**). Analysis using the HDP method also returned the first three mutational signatures (1000G-Sigs 1-3) with cosine similarities >0.93 (**Supplementary Fig. S5**). This method was unable to detect 1000G-Sig4. Altogether, the results obtained from these four independent analyses confirmed the existence of at least three mutational signatures in the 1000 Genomes Project dataset.

**Fig. 2B** shows that Africans have the lowest level of 1000G-Sig1 (*p*<0.0001 pairwise Mann-Whitney U test Bonferroni-corrected) and the highest level of 1000G-Sig4 (*p*<0.0001). The signature 1000G-Sig2 has the highest level in Europeans (*p*<0.0001). Compared to other three signatures, there is less between-population differences in the level of 1000G-Sig3. Nevertheless, in some individuals, except South Asians, the proportion of this mutational signature is over two-fold higher than average (**Fig. 2B**).

**Fig 2.**
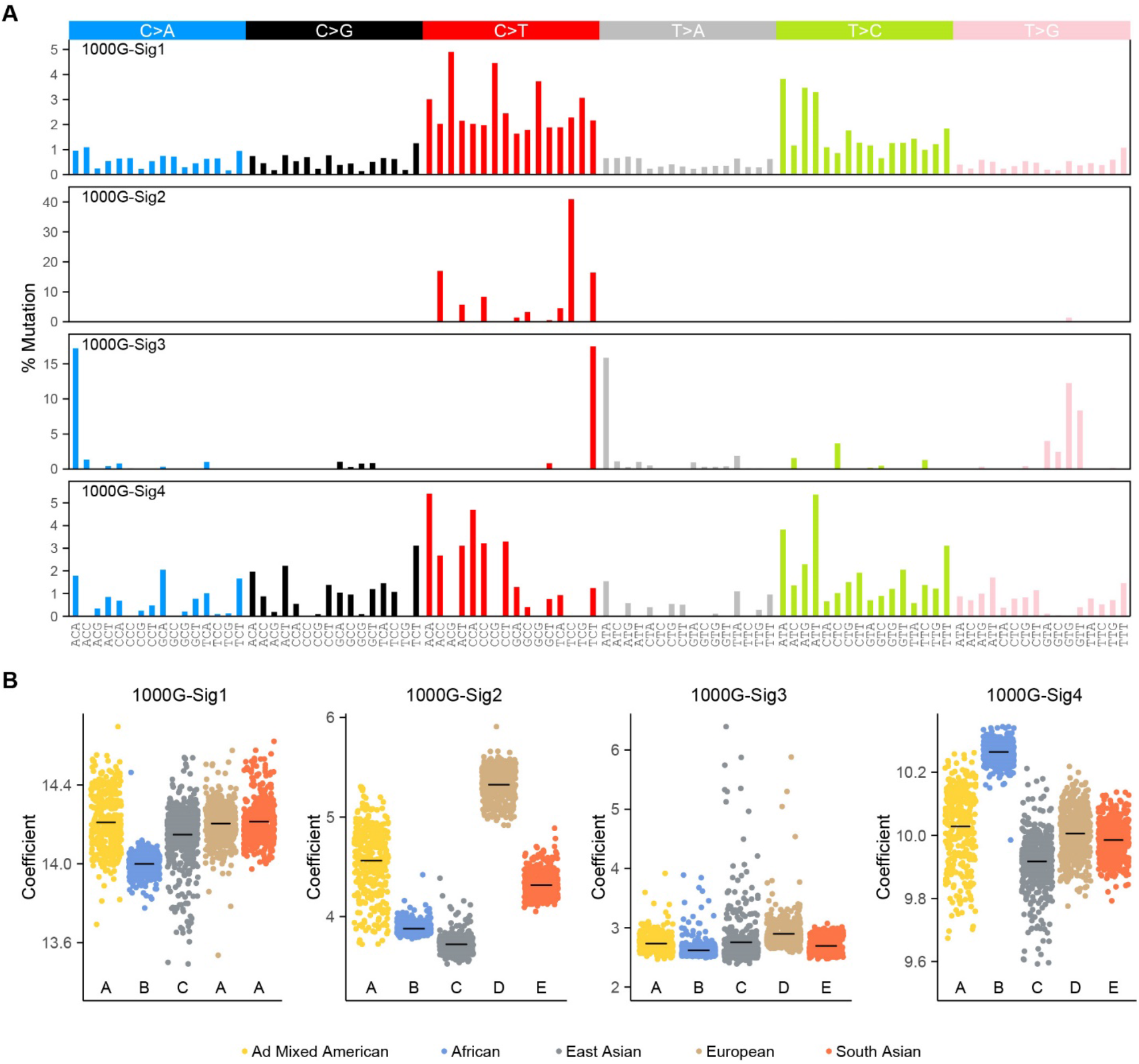
Deconvolution of nuclear DNA mutational signatures in the 1000 Genomes Project dataset using NPCA. (A) Mutational signature 1-4; (B) Coefficient (i.e. weight) of mutational signatures in different populations. Significantly different populations with a Bonferroni-corrected (p<0.01) are denoted by different letters above the horizontal axis.

#### Trio Datasets

Analysis of *de novo* mutations from three trio datasets^20,29,30^ using NPCA showed only a single mutational signature (DN-Sig, **Fig 3**). Same results were found using PCA, NMF, and HDP (**Supplementary Figs. S6, S7, and S8**). This *de novo* mutational signature is almost identical (cosine similarity = 0.997) to the mutational signature 1000G-Sig1.

**Fig 3.**
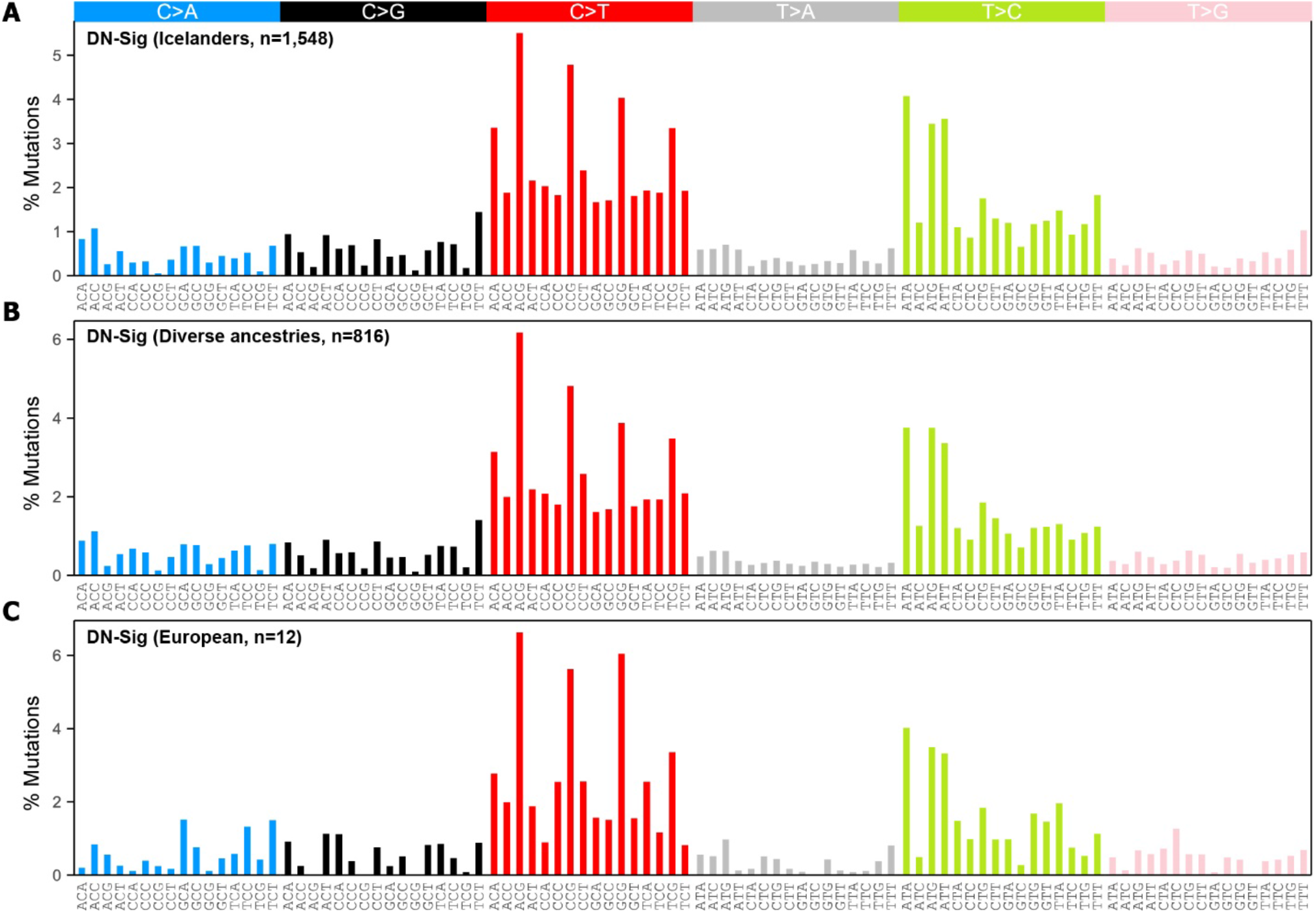
Deconvolution of nuclear DNA *de novo* mutations from two European trio datasets and one dataset containing samples with different ancestry using NPCA. (A) A single mutational signature in Icelanders dataset^13,29^; (B) A single mutational signature in a dataset containing donors from different populations^30^; (C) A single mutational signature in data from Rahbari *et al*.^20^

### Analysis of mitochondrial DNA

#### 1000 Genomes Project Dataset

Typically, in *de novo* mutation studies, mitochondrial DNA and Y chromosome are excluded from the analyses^13,20^. To investigate if, compared to other chromosomes, chrY and mtDNA have a different mutation spectrum, we separated the 1000 Genomes Project mutation datasets by chromosome and computed cosine similarities between each two chromosomes and also between all chromosomes and mtDNA (**Fig. 4A**). The results indicate that all nuclear chromosomes have comparable mutation profiles; however, mtDNA is distinct. Analysis of mtDNA mutations using PCA, NPCA, and NMF returned a single mutational signature (**Supplementary Figs. S9, S10, and S11**). This signature (1000G-Sig-mt) is dominated by C>T and T>C mutations; however, compared to nuclear DNA, there is significantly less CG>TG and TG>CG mutations (**Fig. 4B**). To investigate if this is due to lower representation of CG and TG in mtDNA compared to nuclear DNA, we performed a motif representation analysis. As indicated, compared to nuclear DNA, mtDNA has a significantly higher level of CG and lower level of TG (**Fig. 4C**).

**Fig 4.**
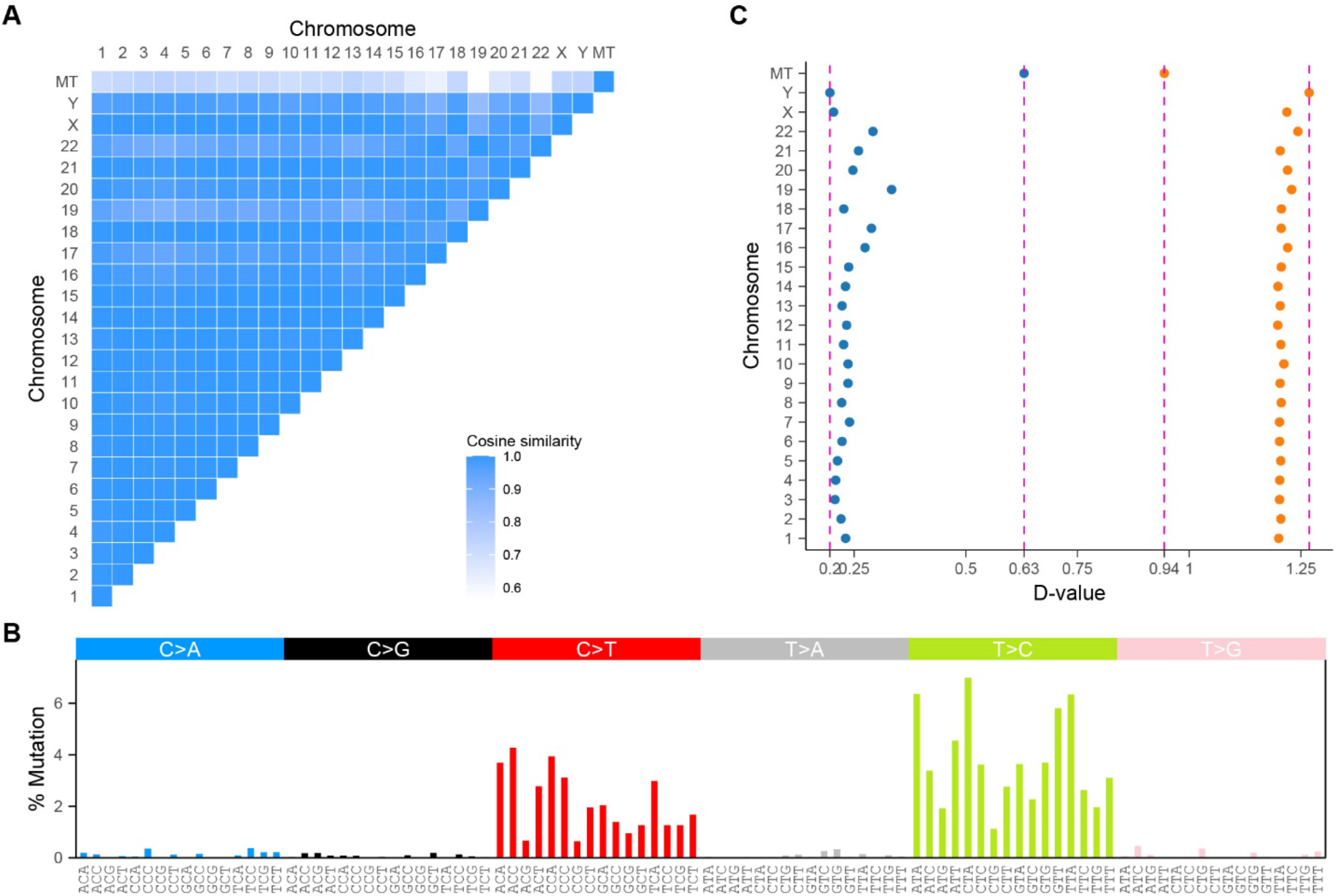
Analysis of mitochondrial DNA in the 1000 Genomes Project dataset using NPCA. (A) Heatmap of similarities between the mutation spectra of chromosomes and mitochondrial DNA; (B) A single mutational signature present in mtDNA (1000G-Sig-mt); (C) Represntation of CG and TG dinucleotides in chromosomes and mitochondrial DNA.

#### Trio Dataset

The number of *de novo* mutations in mitochondrial DNA is very limited (average of ∼18 mutations per donor). Using the available data (three European trios)^20^, it was not possible to perform a deconvolution analysis. However, a sum of mutations in this trio dataset shows a cosine similarity of 0.78 to the 1000G-Sig-mt (**Supplementary Fig. S12**)

### Analysis of similarities between germline and somatic mutational signatures

When compared, to the COSMIC v3 cancer mutational signatures^11,31^, each of the five 1000G Signatures (4 nuclear DNA signatures and one mtDNA signature) and *de novo* signature (DN-Sig). For each germline mutational signature, we observed a cosine similarity of >0.8 to at least one of the recently updated cancer mutational signatures (COSMIC v3) (**Supplementary Table S1**). The 1000G-Sigs 1-4 and 1000G-Sig-mt show the greatest correlations with cancer signatures SBS5, SBS7a, SBS51, SBS5, and SBS12, respectively (**Supplementary Table S1**). Using the expectation maximization (EM) algorithm (Method) we decomposed each of the 1000G-Sigs1,2,3,4,mt signatures into combinations of cancer signatures (COSMIC v3) (**Fig. 5**). Table 1 shows the optimum combinations after the removal of cancer signatures with <10% contribution (Method). The optimum combinations for 1000G-Sigs1,2,3,4,mt are SBS5+SBS1 (**Supplementary Fig. S13**), SBS11+SBS7a (**Supplementary Fig. S14**), SBS47+SBS43+SBS51 (**Supplementary Fig. S15**), SBS39+SBS16+SBS37 (**Supplementary Fig. S16**), and SBS12+SBS30 (**Supplementary Fig. S17**), respectively (**Table 1**).

**Table 1.** Combinations of cancer signatures (COSMIC v3) best describing the 1000G and *de novo* mutational signatures

| COSMIC V3 combination | Original Sigs | Cosine Similarity |
| --- | --- | --- |
| 0.13*SBS1+0.87*SBS5 | 1000G-Sig1 | 1 |
| 0.36*SBS7a+0.64*SBS11 | 1000G-Sig2 | 0.9 |
| 0.26*SBS43+0.58*SBS47+0.16*SBS51 | 1000G-Sig3 | 0.83 |
| 0.29*SBS16+0.23*SBS37+0.48*SBS39 | 1000G-Sig4 | 0.75 |
| 0.68*SBS12+0.32*SBS30 | 1000G-Sig-mt | 0.92 |
| 0.15*SBS1+0.85*SBS5 | DN-Sig | 0.99 |

**Fig 5.**
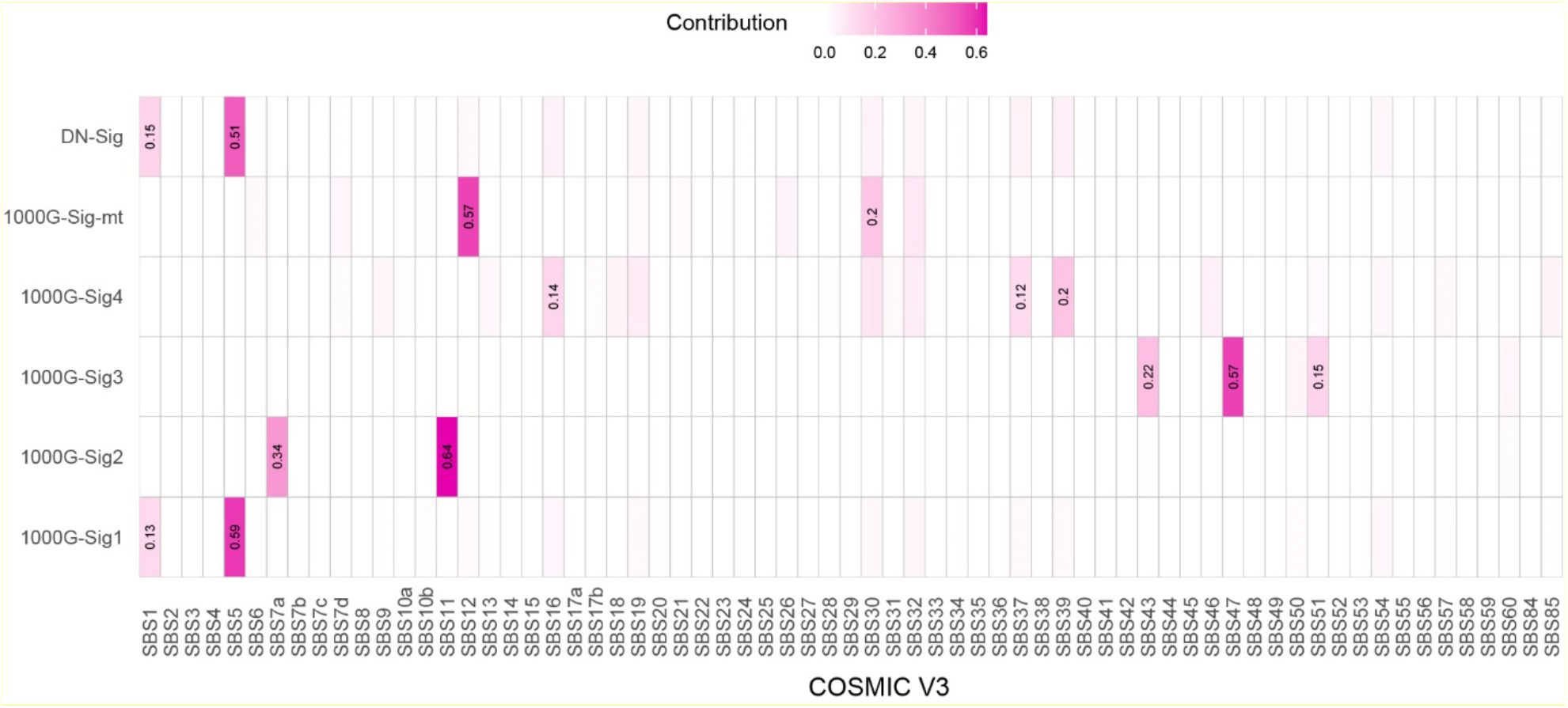
Comparison of germline and COSMIC cancer mutational signatures. The expectation maximization (EM) algorithm was used to determine the combinations of cancer reference signatures (COSMIC v3) that best model each of the five 1000G signatures (1-4 and MT) and DN-Sig. Color intensities in this heatmap represent the contribution of cancer signatures in each germline signature.

Table 1 shows high cosine similarities between 1000G-Sigs and combinations of cancer signatures. Therefore, we investigated if such high similarities could be achieved when distinct cancer mutational signatures (individually or in combination) are compared to each other. Our results showed that cosine similarities between cancer mutational signatures can be as high as 0.99 (**Supplementary Table S2**).

### Enrichment analysis of *de novo* mutations within endogenous elements

Studies suggest that processes such as CpG methylation^32–34^ and APOBEC-induced cytosine deamination^35–37^ play a role in silencing endogenous retroelements. To determine if such processes leave a signature on these elements, we performed an enrichment analysis. We quantified the percentage of mutations occurring within different repeat element (LINEs, SINEs, ERVs, and DNA) and non-repeat regions (e.g. genic regions) for all 1,548 Icelander samples. We first quantified the percentage length of each repeat element class in the human genome to use as a benchmark for mutation enrichment. If *de novo* mutations are enriched within a certain element class, we expect the percentage mutation to be significantly higher than the percentage length of that element. As indicated in **Fig. 6**, the percentage of *de novo* mutations in different regions of the human genome is proportional to the percentage length of the regions. In other words, *de novo* mutations are not enriched within certain repeat element classes (Pearson correlation coefficient = 0.998, p<2.2e-16). However, we note that *de novo* mutations are slightly depleted in the centroid satellite sequences.

**Fig 6.**
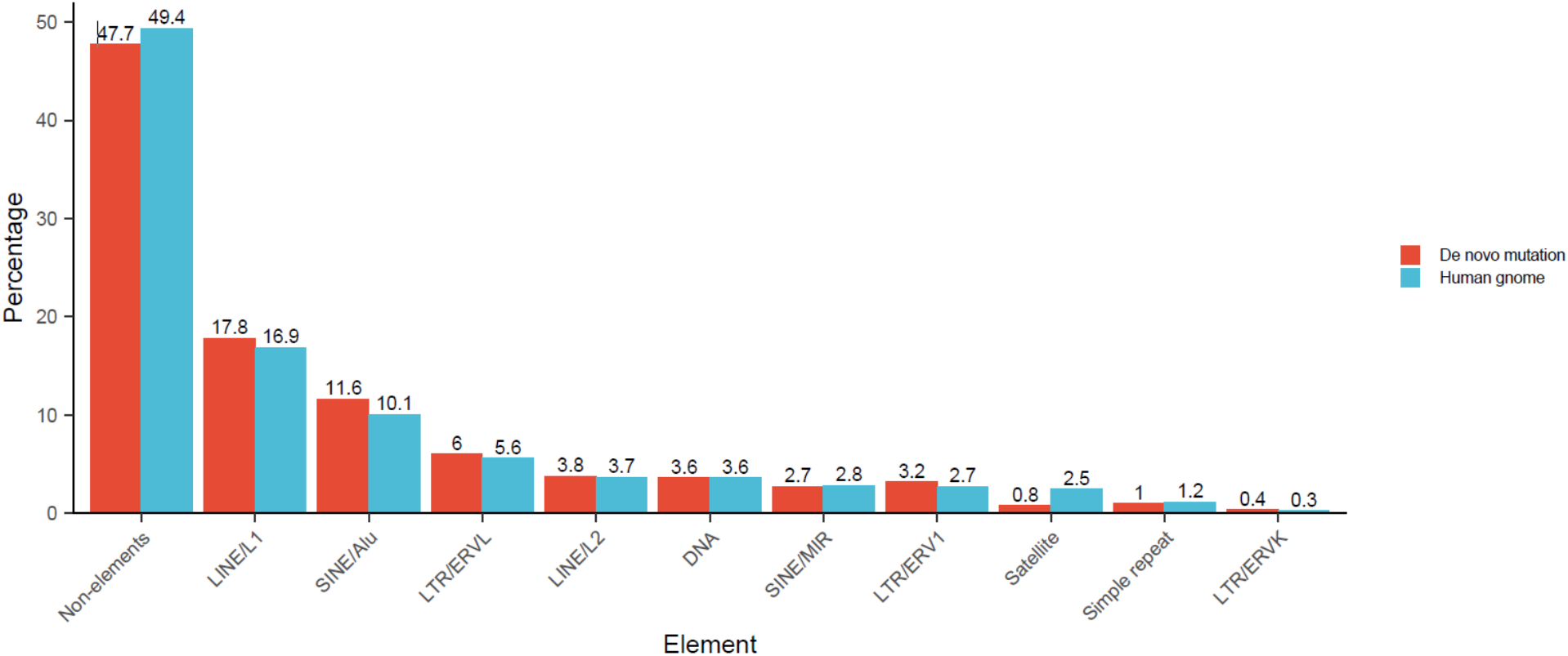
Analysis of *de novo* mutation enrichment within endogenous elements. Percentage of *de novo* mutation within different endogenous elements is shown in red. The total percentage length of each element is shown in blue. Ychr and mtDNA were excluded.

## Discussion

Identification of germline mutational signatures is the first step to discovering the molecular processes responsible for human DNA evolution, their differential triggers in different human populations, and their potential roles in diseases such as cancer. By deconvoluting human genetic variation datasets, we uncovered the mutational signatures imprinted in the human genome by present and past mutational processes. Our analyses show only a single mutational signature (DN-Sig) in germline *de novo* mutation datasets^20,29,30^ This signature is population-independent and likely represents two processes occurring concurrently during gametogenesis and/or early embryogenesis. As a result, their mutational signatures are inseparable by deconvolution methods. One of these processes inflicts C>T within CG (a.k.a. CpG) dinucleotides^38–44^ and the other induces T>C within mostly ATA, ATG, and ATT trinucleotide motifs^20^. The former pattern is identical to the cancer mutational signature SBS1^10,11,31^ and represents methylation-induced deamination of CpG cytosines^21,46,47^. The latter closely resembles SBS5 whose etiology remains unknown^10,11^. SBS1 and SBS5 have also been identified in normal cells^48^ and they show a clock-like behavior happening spontaneously throughout life^10,11,45^. Our data indicate that the CG depletion caused by *de novo* C>T mutations is not fully compensated for by *de novo* T>C mutations, which mostly occur at non-TG motifs. Specifically, at each birth, on average, ∼10 CGs (a.k.a. CpGs) are removed from and ∼4 CGs are introduced to the human nuclear DNA by C>T and T>C mutations, respectively. Assuming that mechanisms such as GC-biased gene conversion^49^ have negligible compensatory impacts, these data suggest that human genome will continue to lose ∼6 CG dinucleotides per birth.

*De novo* mutations are signatures of present-day mutational processes, thus cannot be used to investigate the mutational processes acting in the past. To overcome this limitation, we analyzed the human genetic variation data reported by the 1000 Genomes Project^3,4,8,17^ and identified four distinct mutational signatures in nuclear DNA, and one in mitochondria. The first signature (1000G-Sig1) is almost identical to the *de novo* mutational signature, implying that one can use the 1000 Genomes SNPs to infer *de novo* mutation signatures, without analyzing trio datasets. The proportion of this signature is slightly lower in Africans compared to other populations. The second signature (1000G-Sig2) is composed, almost exclusively, of highly context-specific C>T mutations (TCC>TTC, TCT>TTT, ACC>ATC, and CCC>CTC) and is significantly more prevalent in European populations. Previous studies have identified most of these trinucleotide motifs within which C>T changes are enriched in Europeans^3–5,17,28^. Our matrix deconvolution approaches precisely identified these mutation types and grouped them into a distinct mutational signature. Our analysis of present-day trios did not show this mutational signature, confirming that 1000G-Sig2 has an ancient source^3,4,27,28^. The signature 1000G-Sig2 can be modelled by a linear combination of two cancer signatures SBS7a and SBS11. The former is associated with skin cancer and exposure to UV light, and the latter resembles the mutations caused by alkylating agents. UV-mediated degradation of folate leading to defects in DNA synthesis were suggested to be a potential source of germline mutations in Europeans with light skins^3^. Mathieson *et al*. argued against this hypothesis based on two main lines of evidence^17^. First, ACC>ATC is not enriched in the genomes of other lightly pigmented populations such as Siberians and Northeast Asians. Importantly, our data also do not show an enrichment for 1000G-Sig2 in East Asians. Second, UV irradiation is known to induce CC>TT mutations, but this type of mutation is not enriched in West Eurasians^17^. With respect to alkylating agents, there is no evidence to suggest a historical exposure of Europeans to these chemicals, although it is suggested by Mathieson *et al*.^17^ Taken together, a high similarity between 1000G-Sig2 and SBS7a+SBS11 may not be sufficient evidence to suggest exposure to UV radiation and alkylating agents have played a role in human genome diversification.

All of the three cancer signatures SBS47, SBS43, and SBS51 predicted to be related to 1000G-Sig3 are currently deemed to be sequencing artefacts. Additionally, this signature contains all of the four peaks of GTN>GGN (NAC>NCC), which were first reported as a Japanese-specific signature^4,5^, but later found to be a batch effect^50^. These data suggest our analyses may have resolved a systematic sequencing error present in the 1000Genome Project dataset. The mutational signature 1000G-Sig4 may also be an artefact of the analysis. This signature was found by PCA, NPCA, and NMF, but not HDP. Additionally, compared to other signatures, it shows the lowest coefficients and model accuracy when reconstructed using a combination of cancer signatures. These suggest further analyses are needed to determine if this signature represents a real mutational process. Nevertheless, 1000G-Sig4 exhibits a T>C profile that resembles SBS16 (and SBS5), and a C>T profile that is highly depleted for CG>TG mutations. This is the opposite of *de novo* signature DN-Sig in which T>C mutations are always accompanied by C>T mutations at CG sites. Assuming that 1000G-Sig4 is not an artefact, these data suggest that T>C mutations might have happened independently of CG>TG mutations in the past, particularly in Africans with an elevated level of 1000G-Sig4, but these two mutation types co-occur in the present time.

We found that compared to nuclear DNA, mtDNA has a distinct mutational signature featured by C>T changes at mostly non-CG sites and T>C changes at mostly non-TG sites. To investigate if the lower rates of CG>TG and TG>CG are due to the low abundance of CG and TG in the mtDNA compared to nuclear DNA, we performed a motif representation analysis. As indicated in, TG dinucleotide is ∼3 folds less abundant in mtDNA, suggesting that lower frequency of TG in mtDNA may be a player in reduced TG>CG level. Nevertheless, further studies are needed to confirm this hypothesis. As opposed to TG, the representation of CG is significantly higher in mtDNA (∼4 folds), implying that the low rate of CG>TG mutation in mtDNA is due to another mechanism. One hypothesis could be that compared to nuclear DNA, the rate of CG methylation (followed by spontaneous deamination) is lower in mtDNA. In support of this hypothesis, a recent study by Patil *et. al*. provides evidence that methylation of human mtDNA mostly occurs at non-CG sites^51^. Importantly, the majority of somatic mutations identified in mtDNA from tumor samples are also predominantly C>T and T>C changes^52^. But the sequence contexts of these somatic mutations are very different from germline *de novo* mutations. For example, contrary to *de novo* germline mutations, which are depleted for CG>TG mutations, these are the most frequent mtDNA mutations in cancer^52^. These data suggest that the etiologies of germline and somatic mtDNA mutations are different.

Silencing of retroelements using mechanisms such as methylation and APOBEC-induced deamination are well documented^32–37,53,54^. Methylation-induced deamination of cytosines in these elements during gametogenesis, fertilization and/or in early embryogenesis can be a potential source of *de novo* mutations. Such targeted mechanisms would lead to a *de novo* mutation enrichment within retroelements. However, our systematic analysis of mutations in all main classes of endogenous elements did not show signs of mutation enrichment in any element classes. These data suggest that epigenetic silencing of retroelements is not likely a player in the observed pattern of *de novo* mutation.

Altogether, human polymorphism datasets contain at least three mutational signatures that are likely associated with biological processes. One of these signatures is currently active and appears to represent two co-occurring mutational processes, methylation-induced deamination and an unknown process. The second signature, which is elevated in the genome of Europeans, is no longer active^3,4,27^. The third signature is specific to mitochondrial DNA and represents mutational process(es) that inflict C>T and T>C changes but mostly at non-CG and non-TpG sites. Each of these three signatures (and also the other two signatures that might be artefacts) show unexpectedly high cosine similarities with combinations of COSMIC cancer signatures. This may indicate, at first glance, that germline mutations are simply a combination of two or more processes already identified in cancer. Our analyses show that cancer mutational signatures can also be modelled, with cosine similarities of up to 0.99, using combinations of other cancer signatures. This implies a high similarity between a germline signature and a combination of known cancer signatures cannot always be used to infer mutational processes.

## Methods

### Data

In this study we used four independent datasets: a) Phase 3 of the 1000 Genomes Project dataset, which includes whole genome SNPs of 2504 individuals^2^, b) whole-genome *de novo* mutation of 1,548 trios from Iceland^29^, and c) whole genome *de novo* mutation dataset of three multi-sibling families^20^ d) whole-genome *de novo* mutation dataset of 816 trios with different ancestry^30^. For each dataset, we performed a separate analysis using all mutations except those from mtDNA DNA and Y chromosome for which only had access to data from the first and third datasets. For the analysis of mutation distributions, we used the Repeatmasker dataset to determine the location of various DNA elements such as LINE, SINE, ERVs, etc^55^.

### Analysis of 1000 Genomes SNPs

We used the FASTA file of the most common ancestral sequence (MCA) (http://ftp.ensembl.org/pub/release-74/fasta/ancestral_alleles/) as a reference and compared each of the 2,504 individuals to this reference sequence using available VCF files (http://ftp.1000genomes.ebi.ac.uk/vol1/ftp/release/20130502/). Using this method, we identified, for each individual and each polymorphic position, the type of mutations with respect to the ancestral sequence. For example, if a position in the ancestral FASTA file is T, and the studied individual has a G in the same position, there has been a T>G mutation in this position. Considering all possible 3’ and 5’ flanking nucleotides of a mutated base, there would be a total of 192 types of mutations (e.g. AAA>ACA, AAA>AGA, …, TTT>TGT). This number would be halved if we combine each mutation with its reverse complement (e.g. ACT>ATT and AGT>AAT). We identified the trinucleotide context of each of the mutated positions described above and created a matrix of 96 x 2,504. Each cell in this matrix represents the total number of one of the 96 type of mutations in one individual. In our analyses, we did not include the SNPs for which there was no information on the type of nucleotides in the MCA FASTA file. To avoid the potential effect of selection, previous studies have focused on rare variants^17^. Previous studies have also removed singletons, which are mutations occurring in only one sample out of the entire population and might represent cell line and/or somatic mutation artifacts^17^. We removed, from our analyses, all singletons and all SNPs with an allelic frequency >1%. We used four independent analyses, namely PCA, NPCA, NMF, and HDP on the matrix of mutations, which included all chromosomes except chromosomes Y and mitochondria DNA. We performed separate analyses for mtDNA. For parsing the VCF and FASTA files we used the “pyvcf^56^” and “biopython^57^” libraries in Python. We noted that the number of all 96 mutation types were 2-4 folds higher in African populations compared to other populations (**Supplementary Fig. S18**). Therefore, we normalized the data such that within each donor, each mutation is presented as a percentage of total number of mutations. The NPCA, NMF, and HDP analyses were done on this normalized matrix of mutations.

### Analysis of *de novo* mutations in trios

Using the reference parental alleles, the derived alleles of the offspring, and the human reference sequence GRCh38 for Icelandic trios and GRCh37 for 816 trios with different ancestry, we identified the type and sequence contexts of all *de novo* mutations and created a matrix of size (96 mutation type × number of individuals). We then performed NPCA, PCA, and NMF as described below on each dataset. We also analyzed the mixed-matrix of all datasets consisting of 2364 individuals using HDP.

### Deconvolution methods

#### PCA

The PCA analyses were performed using the ‘prcomp’ function in R^58^, which uses a singular value decomposition algorithm.

#### NPCA

The NPCA analyses were performed using the ‘nsprcomp’ package^22^ in R^58^, and only the non-negativity constrained was used to perform PCA. We performed NPCA with 100 random initializations to avoid local minima solutions.

#### NMF

Alexandrov *et al*. developed a six-step mutation deciphering method, which has a non-negative matrix factorization step to identify signatures of mutational processes in cancer^10,16,59^. We used these steps to identify mutational processes in *de novo* mutations from three trio datasets. We used both signature reproducibility and reconstruction error measures to determine the optimum number of mutational signatures. For the analysis of 1000 Genomes dataset, we used the elbow point of explained variance to identify the number of mutational processes^60,61^. We used the “NMF” package^60^ in R programming language^58^ with 100 random initialization (to avoid local minima) and “brunet” algorithm^62^ to perform NMF. We noted that the NMF signature 1 peaks appear as background in all other three signatures 2, 3, and 4. To remove this background, we subtracted NMF signatures 2, 3, and 4 from NMF signature 1, and then determined the cosine similarities between this background-corrected NMF signatures and NPCA signatures.

#### HDP

To extract mutational signatures, we ran an algorithm based on the hierarchical Dirichlet process (HDP)^23^ (https://github.com/nicolaroberts/hdp) on the 96 trinucleotide counts. HDP was run with individuals or - where available - the population as the hierarchy, in twenty independent chains, for 40,000 iterations, with a burn-in of 20,000.

### Signature reconstruction using COSMIC reference signatures

The resulting 1000G-Sigs were decomposed into optimal combinations of cancer signatures (COSMIC v3) using expectation maximization (EM) algorithm^24^. First, the EM algorithm was performed on all COSMIC signatures. Afterward, EM algorithm was performed on signatures with contribution of more than 10% to avoid overfitting. The EM algorithm finds the optimal linear combination of chosen COSMIC signatures that reconstructs our 1000G-Sigs. The cosine similarity between reconstructed signatures and original signatures was computed.

### Analysis of motif representation

We observed that the mutation profile of mitochondria DNA was different from that of nuclear DNA. To investigate if this difference is due to the genetic makeup difference between nucleus and mitochondria, we quantified the ratio of observed frequency of each dinucleotide motif over its expected frequency. The observed frequency is, simply, the proportion of a given motif in the DNA. The expected frequency of a dinucleotide motif is calculated by multiplying the constituent mononucleotides^63^.

### Analysis of *de novo* mutation enrichment within DNA elements

We investigated the potential enrichment of mutations within major endogenous elements, LTR (ERV1, ERVL, ERVK), LINE (L1 and L2), SINE (Alu and MIR), DNA, and/or other minor classes. We used Repeatmasker to download the genomic coordinates of these elements, available at (http://www.repeatmasker.org/genomes/hg38/RepeatMasker-rm405-db20140131/hg38.fa.out.gz). We quantified the percentage of the length of each element class in the human reference genome, excluding chrY and mtDNA. We determined the mutation burdens by quantifying the number of substitutions within each element class listed in the repeat masker file, summed over all 1,548 probands. These numbers were divided by the total number of *de novo* mutations to calculate the percentage of mutations within each element class. We used the Pearson’s product moment correlation coefficient in R to investigate the statistical difference.

### Analysis of mitochondrial DNA *de novo* mutations

Mutations were initially called using bcftools mpileup using the 1000Genomes_hs37d5 as reference. Variants with a quality of less than 20 and a depth of greater than 100 reads were excluded. Sequencing of these individuals is described in Rahbari *et al*.^20^

## Acknowledgement

The authors thank Mrs. Rachael Rodriguez for her scientific comments and proofreading the manuscript.

---

**Supplementary Figure S1.**
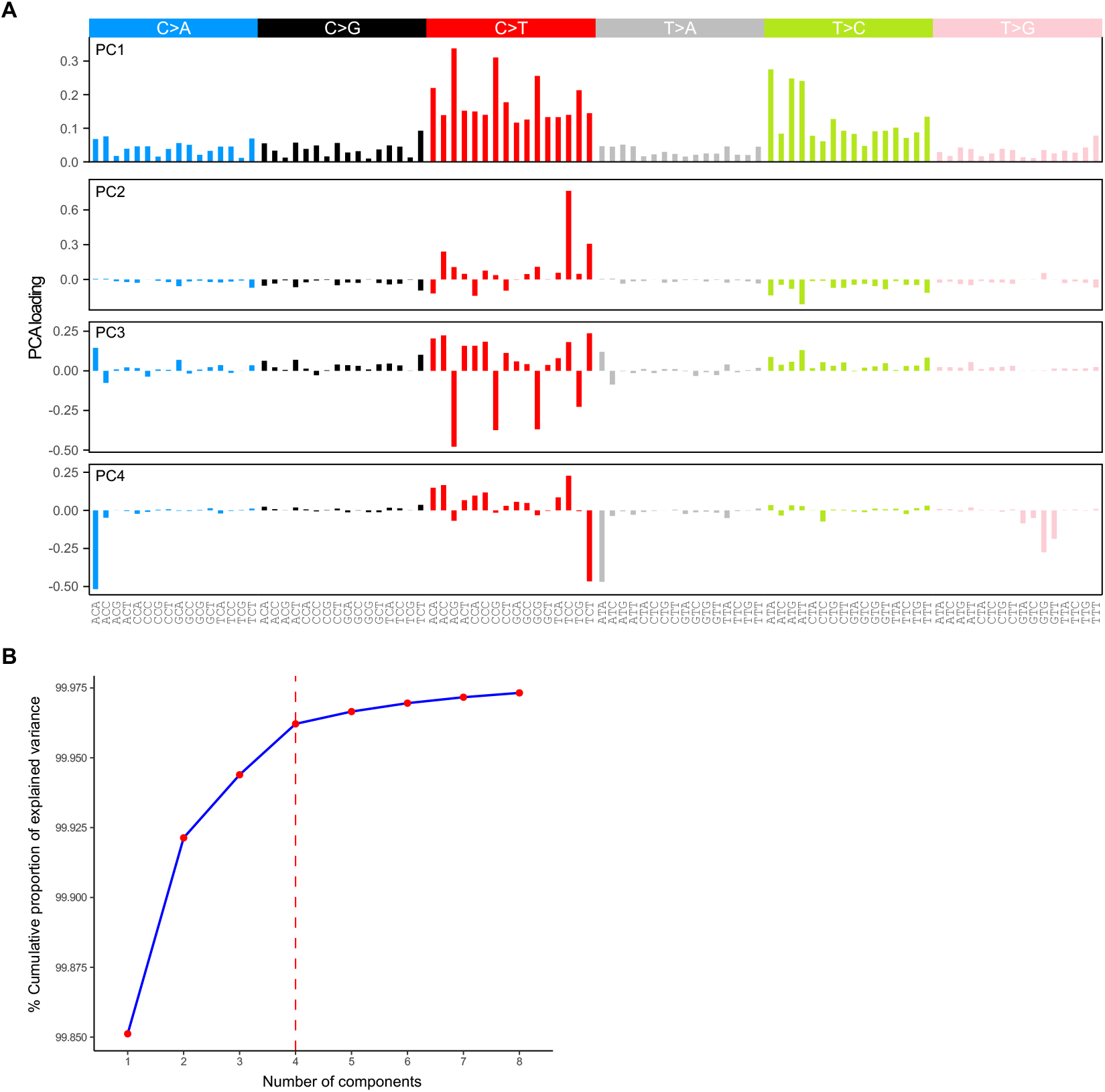
Analysis of the 1000 Genome Project dataset using PCA. (A) First four loadings plots. (B) Evaluation plot showing the %cumulative variance in PCA models with different number of components.

**Supplementary Figure S2.**
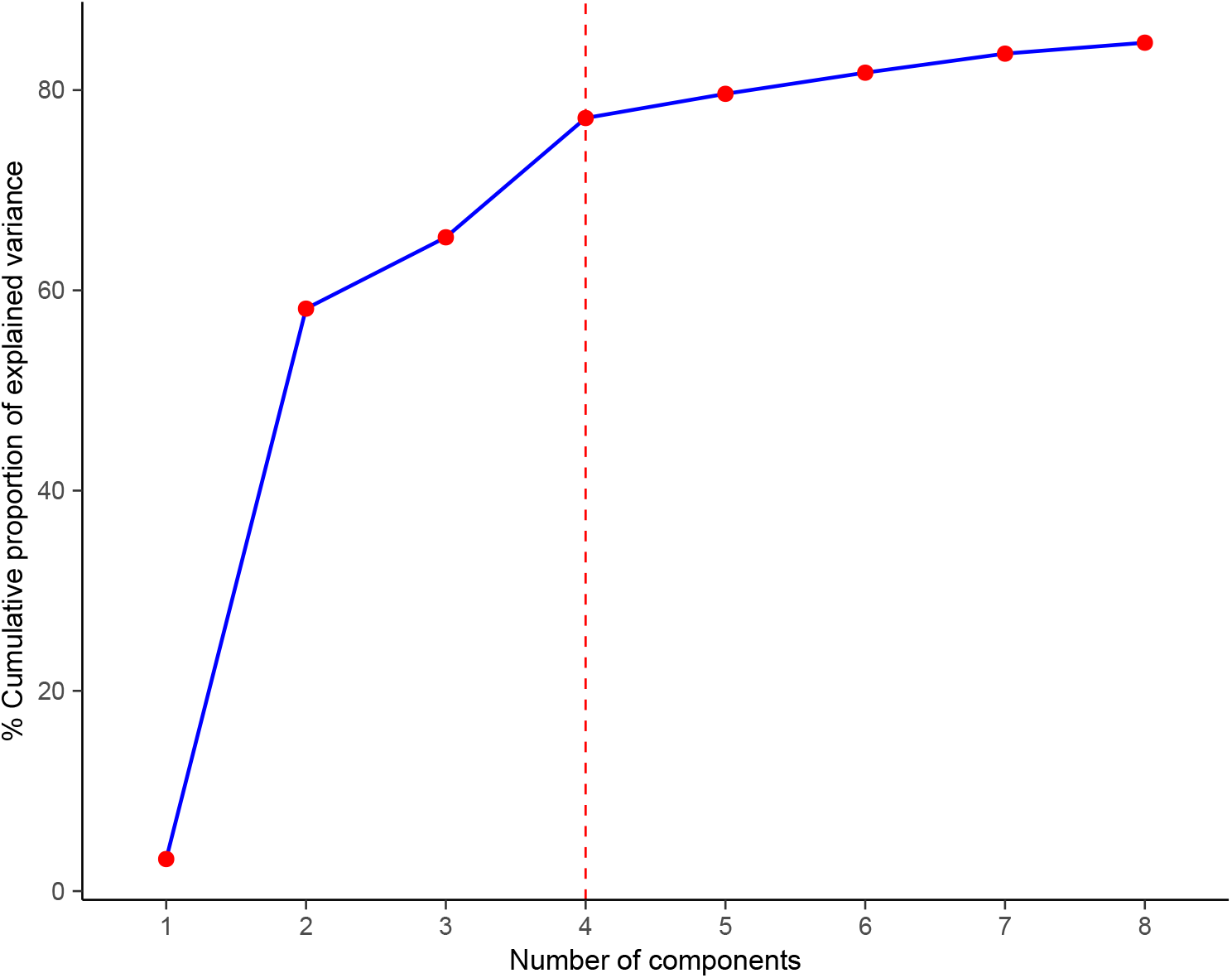
Analysis of the 1000 Genome Project dataset using NPCA. Evaluation plot showing the %cumulative variance in NPCA models with different number of components.

**Supplementary Figure S3.**
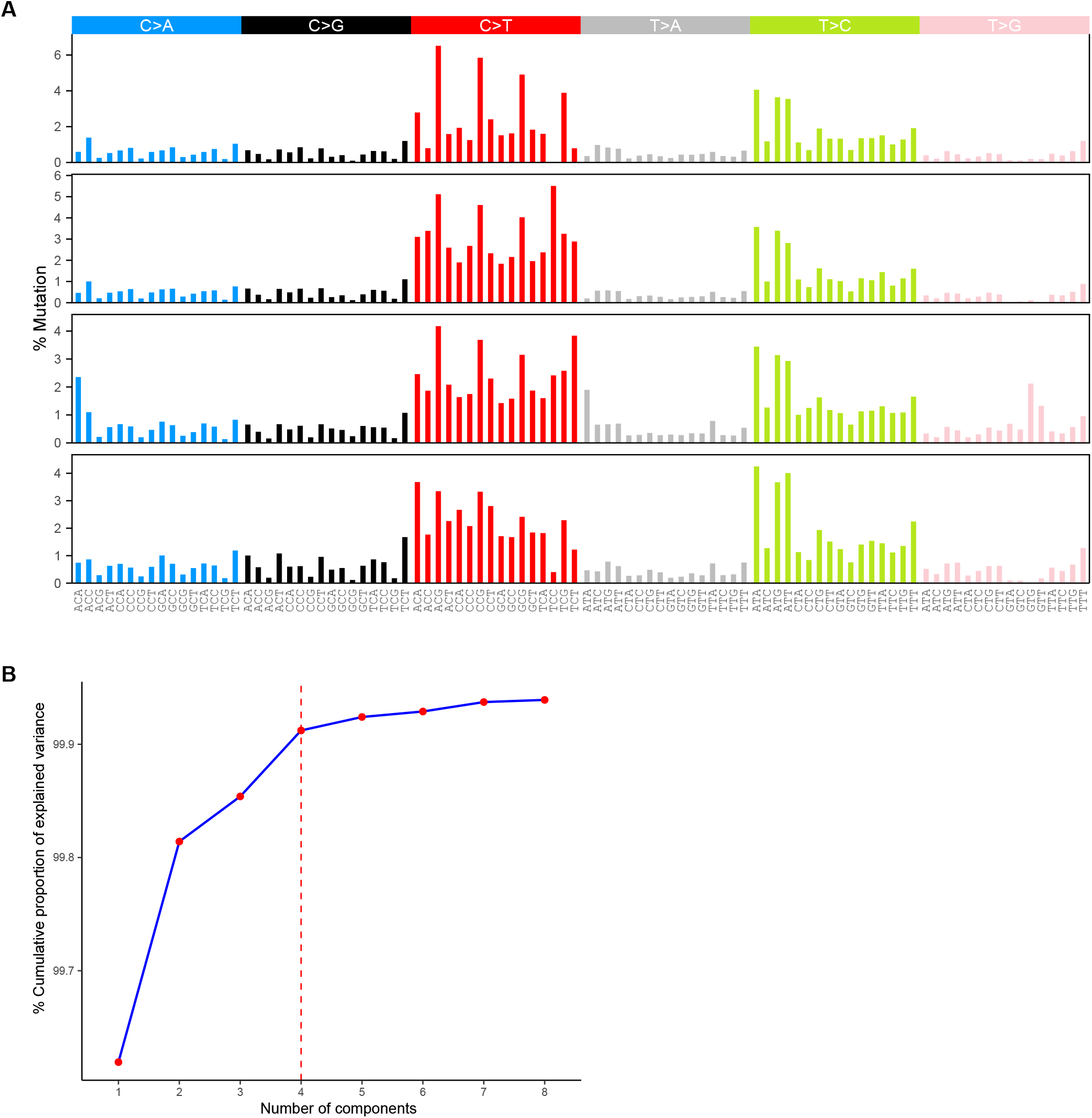
Analysis of the 1000 Genome Project dataset using NMF. (A) Mutational signatures 1-4. (E) Evaluation plot showing the %cumulative variance in NMF models with different number of components.

**Supplementary Figure S4.**
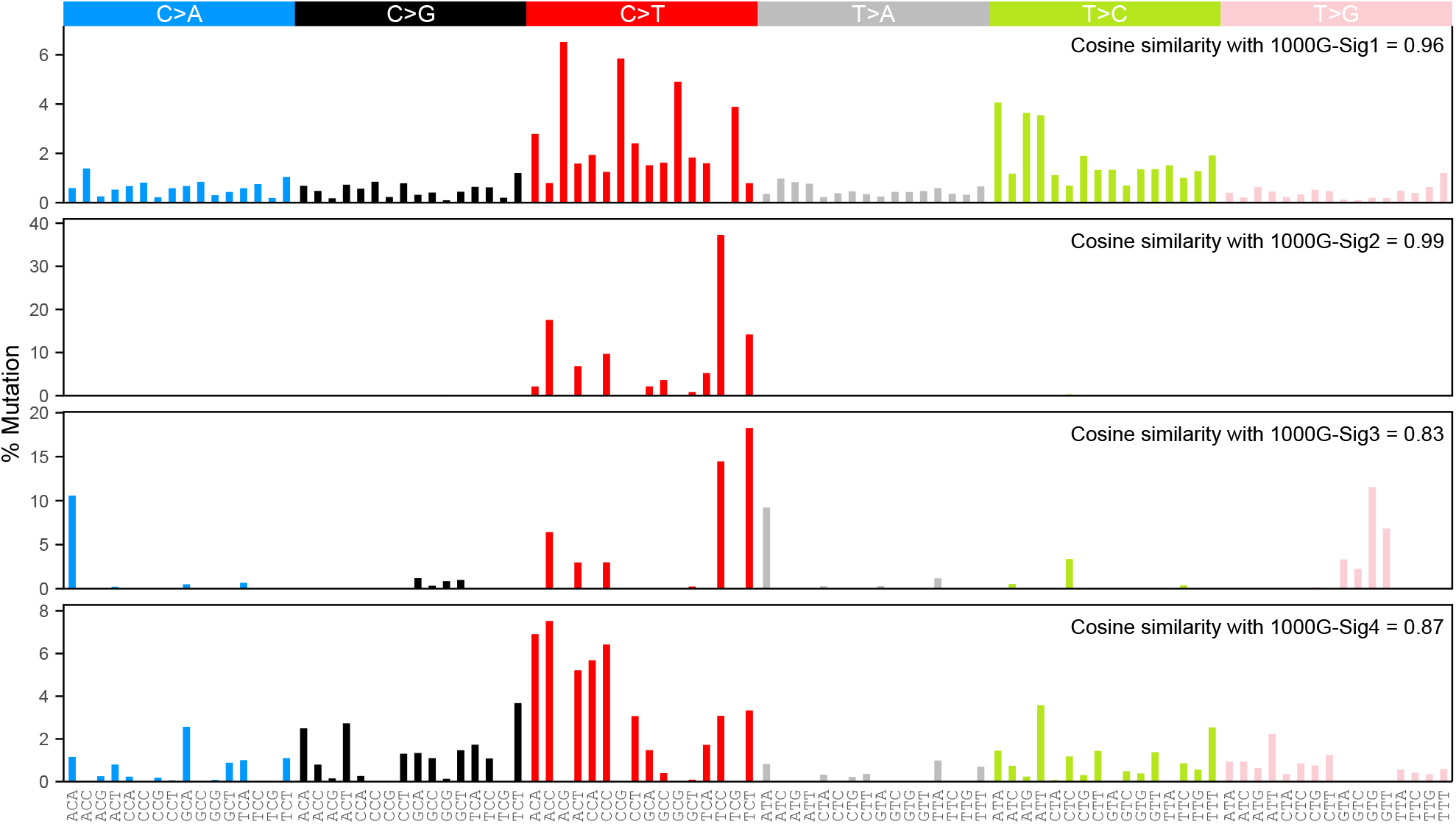
Analysis of the 1000 Genome Project dataset using NMF followed by the removal of a common background from signatures 2-4. NMF signatures 1-4 and their corresponding cosine similarities to the 1000G-Sigs 1-4 obtained using NPCA.

**Supplementary Figure S5.**
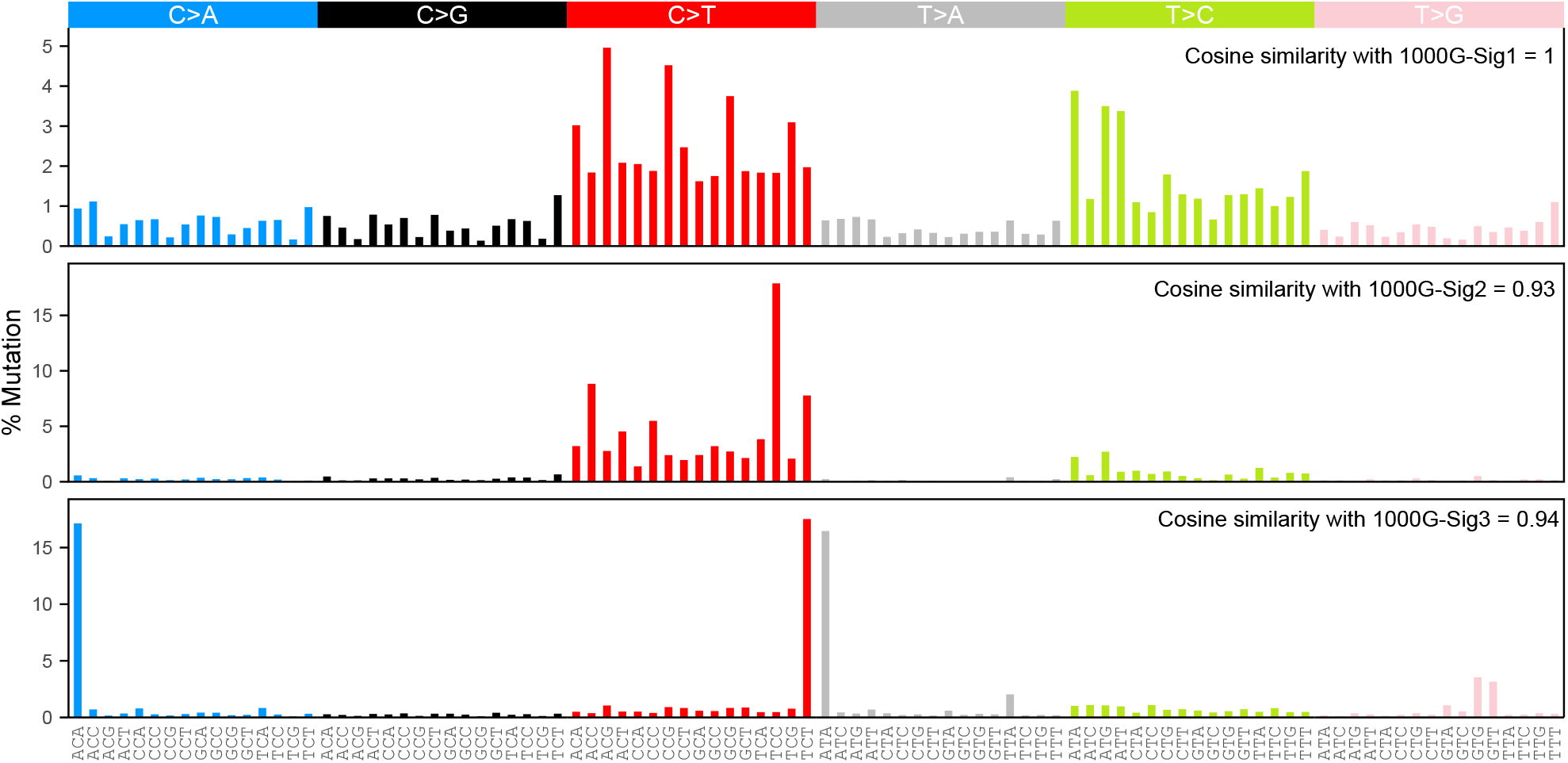
Analysis of the 1000 Genome Project dataset using HDP. Three identified mutational signatures and their corresponding cosine similarities to the 1000G-Sigs 1-3, obtained using NPCA, are shown.

**Supplementary Figure S6.**
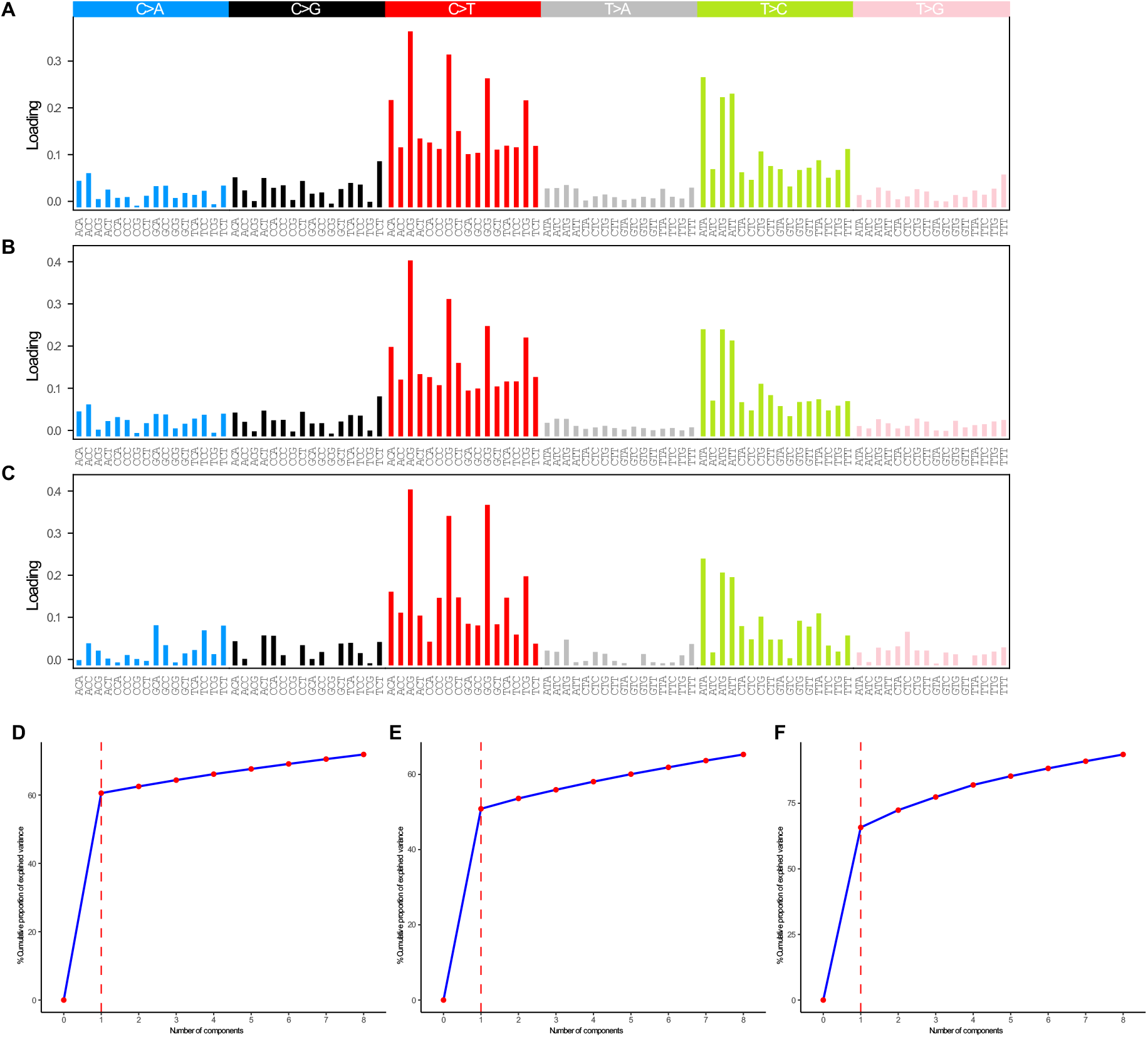
Analysis of three *de novo* mutation datasets using PCA. (A) Mutational signature obtained from the analysis of a dataset of 1,548 Icelander trios. (B) Mutational signature obtained from the analysis of a dataset of 816 trios from diverse populations (dbGap). (C) Mutational signature obtained from the analysis of a dataset of 12 European trios. (D-E) Evaluation plots showing the %cumulative variance in PCA models with different number of components for datasets A, B, and C, respectively.

**Supplementary Figure S7.**
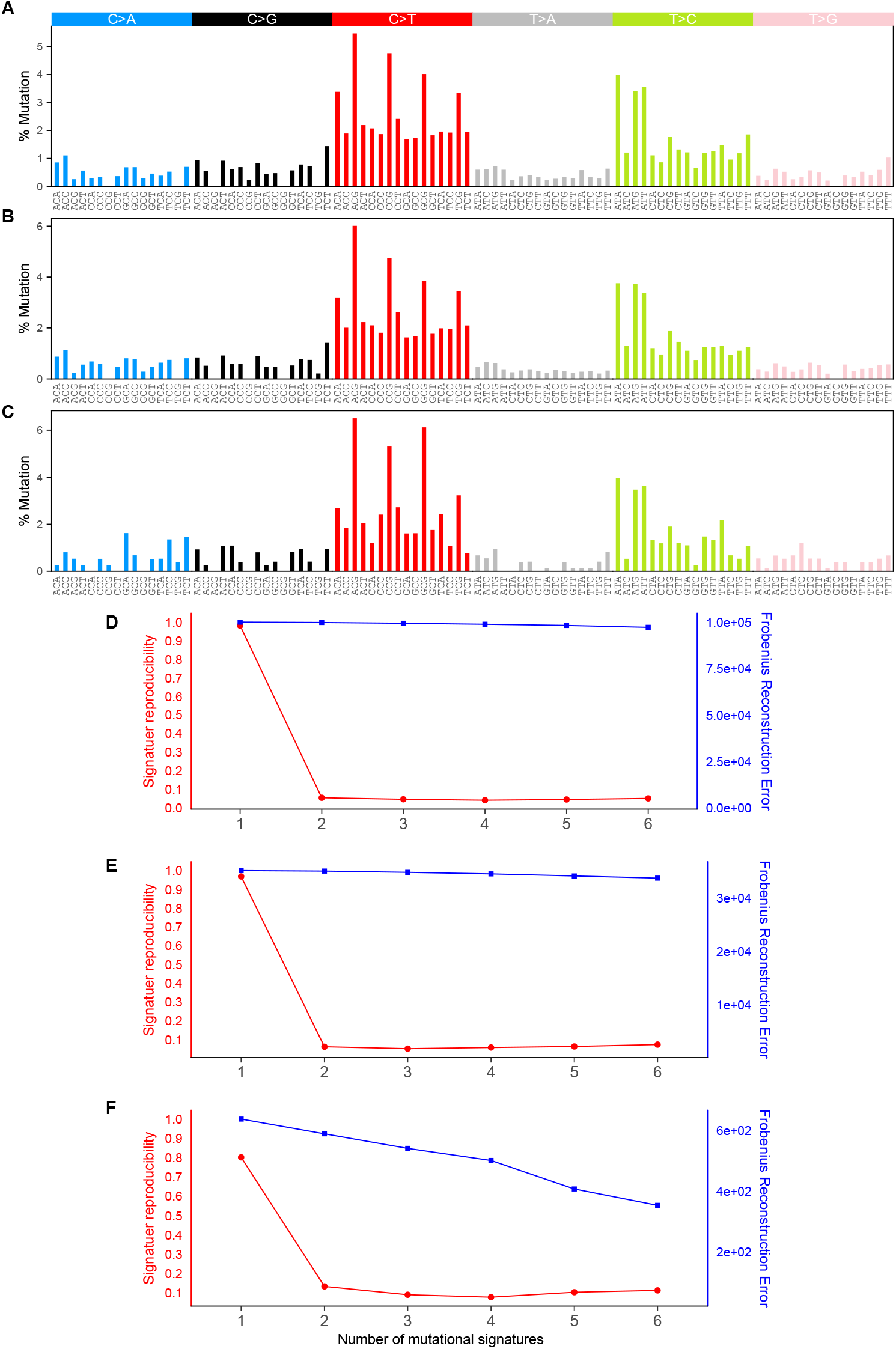
Analysis of three *de novo* mutation datasets using NMF. (A) Mutational signature obtained from the analysis of a dataset of 1,548 Icelander trios. (B) Mutational signature obtained from the analysis of a dataset of 816 trios from diverse populations (dbGap). (C) Mutational signature obtained from the analysis of a dataset of 12 European trios. (D-E) Evaluation plots showing the signature reproducibilities and reconstruction errors in NMF models with different number of components for datasets A, B, and C, respectively.

**Supplementary Figure S8.**
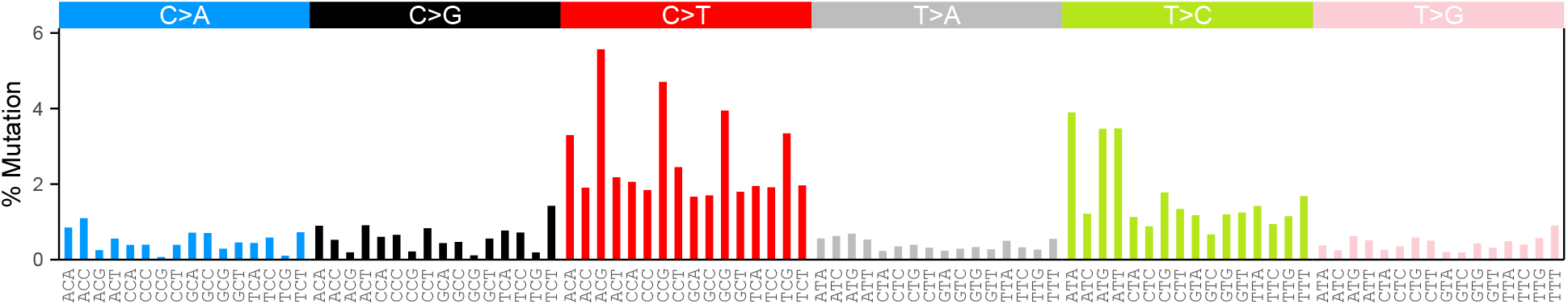
Analysis of three *de novo* mutation datasets using HDP. One mutational signature was identified.

**Supplementary Figure S9.**
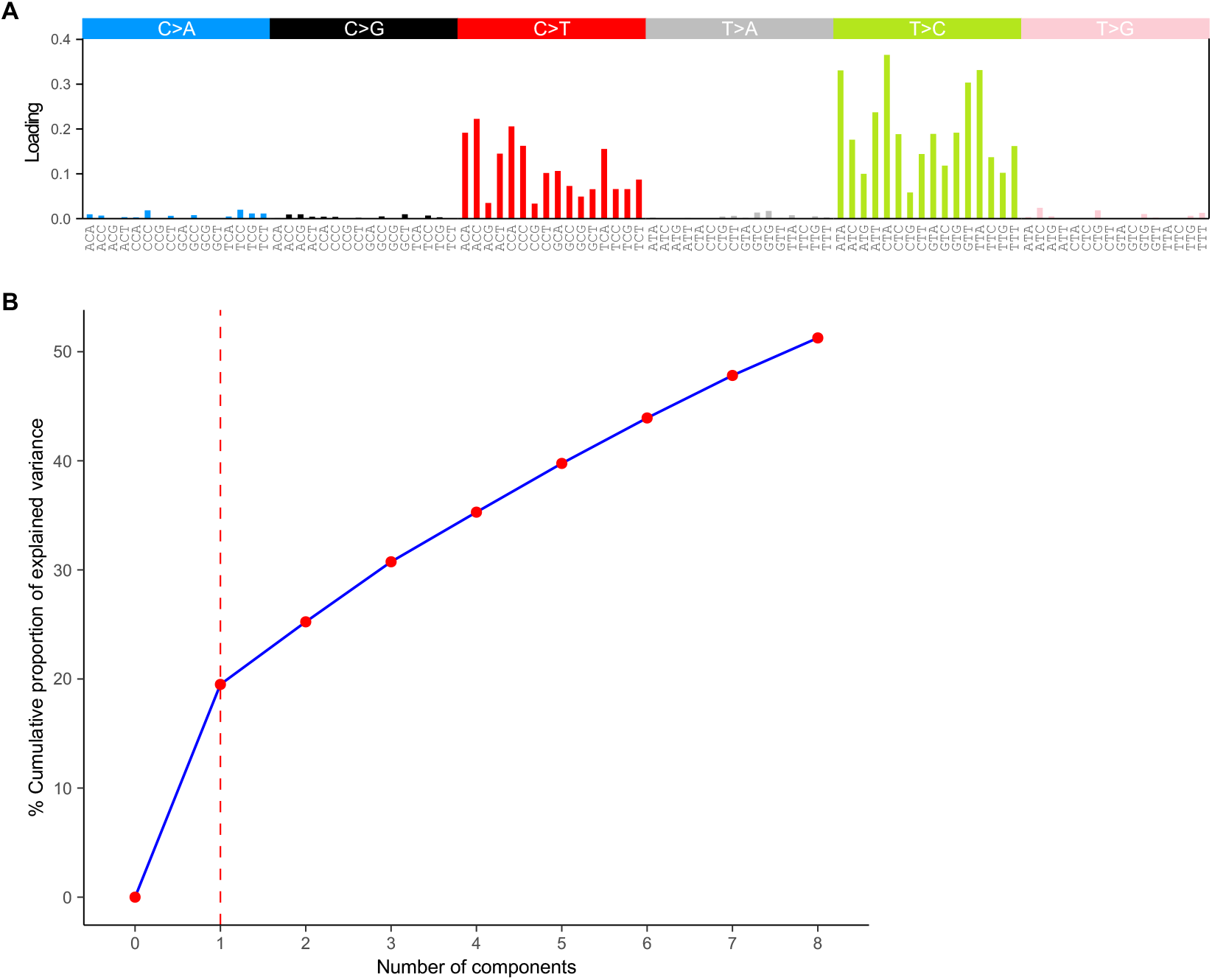
Analysis of mtDNA mutations using PCA. (A) The mtDNA mutational signature obtained using PCA (B) Evaluation plot showing the %cumulative variance in PCA models with different number of components.

**Supplementary Figure S10.**
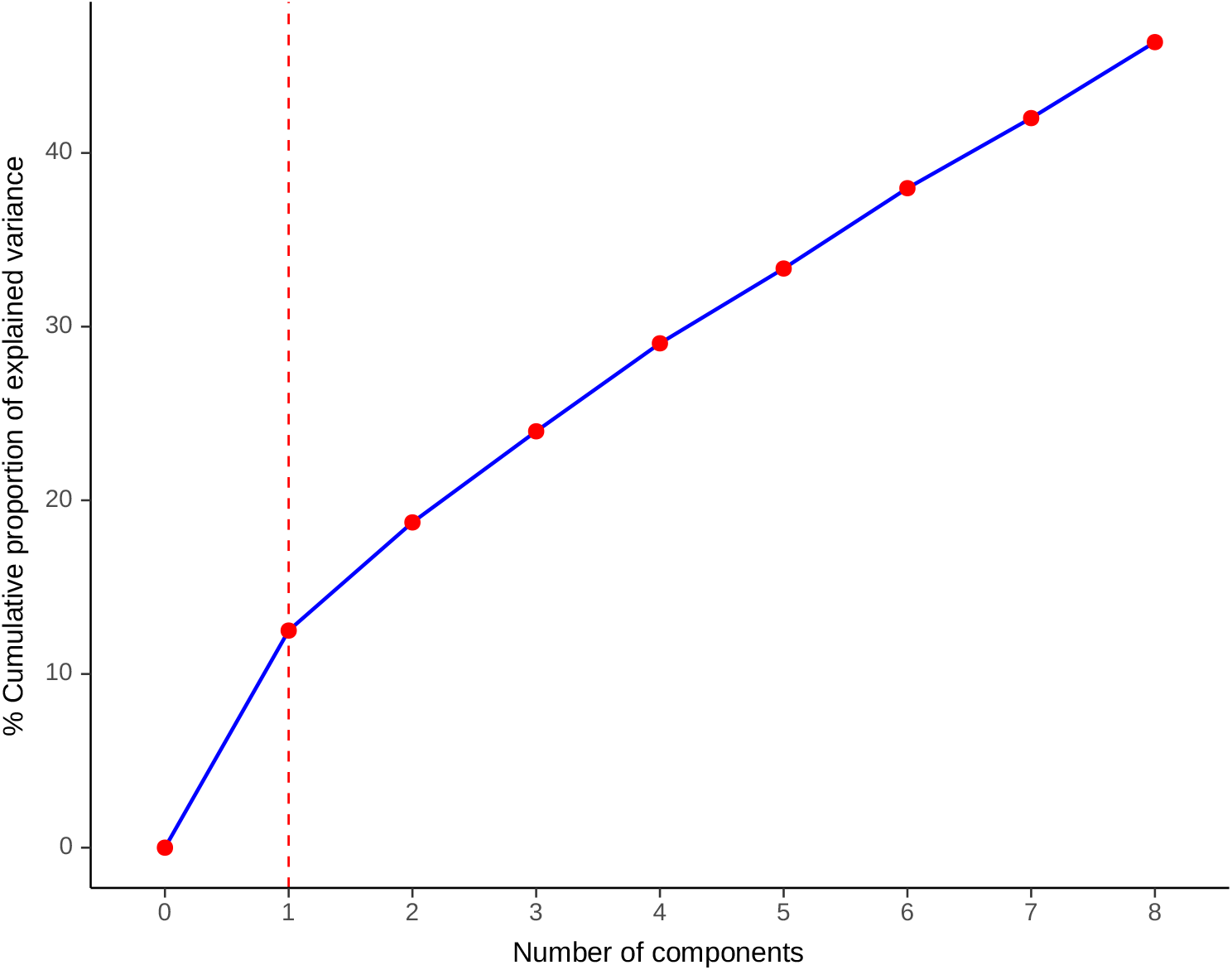
Analysis of mtDNA mutations using NPCA. Evaluation plot showing the %cumulative variance in NPCA models with different number of components.

**Supplementary Figure 11.**
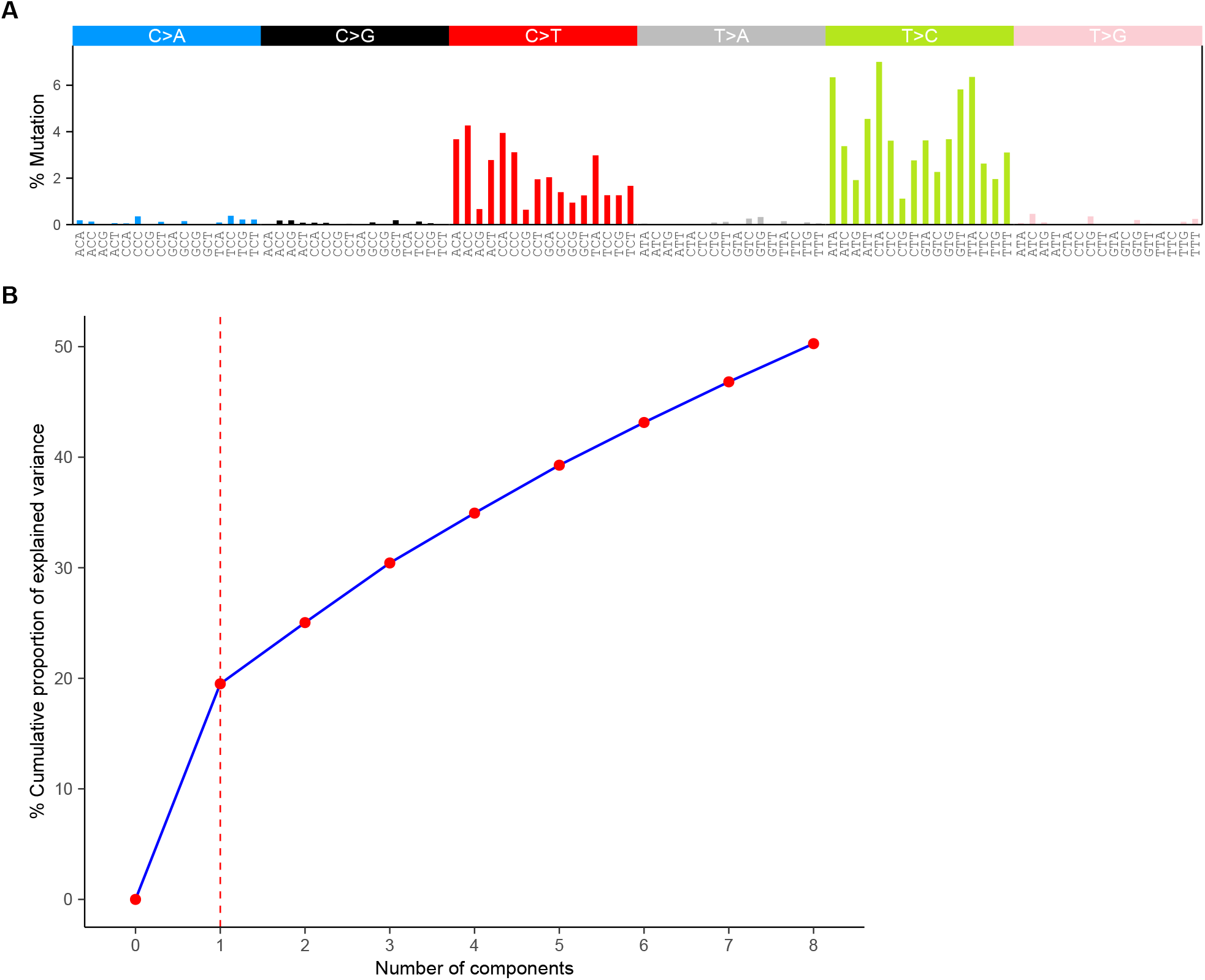
Analysis of mtDNA mutations using NMF. (A) The mtDNA mutational signature obtained using NMF (B) Evaluation plot showing the %cumulative variance in NMF models with different number of components.

**Supplementary Figure S12.**
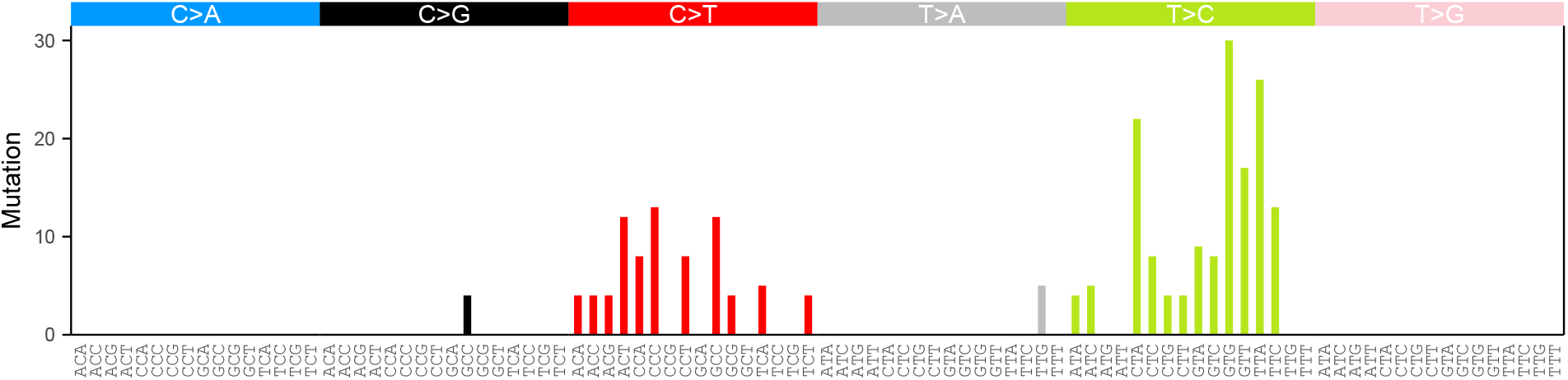
The mutation profile of *de novo* mutations in mitochondrial DNA. The cosine similarity between this profile and 1000G-mt-Sig is 0.78.

**Supplementary Figure 13.**
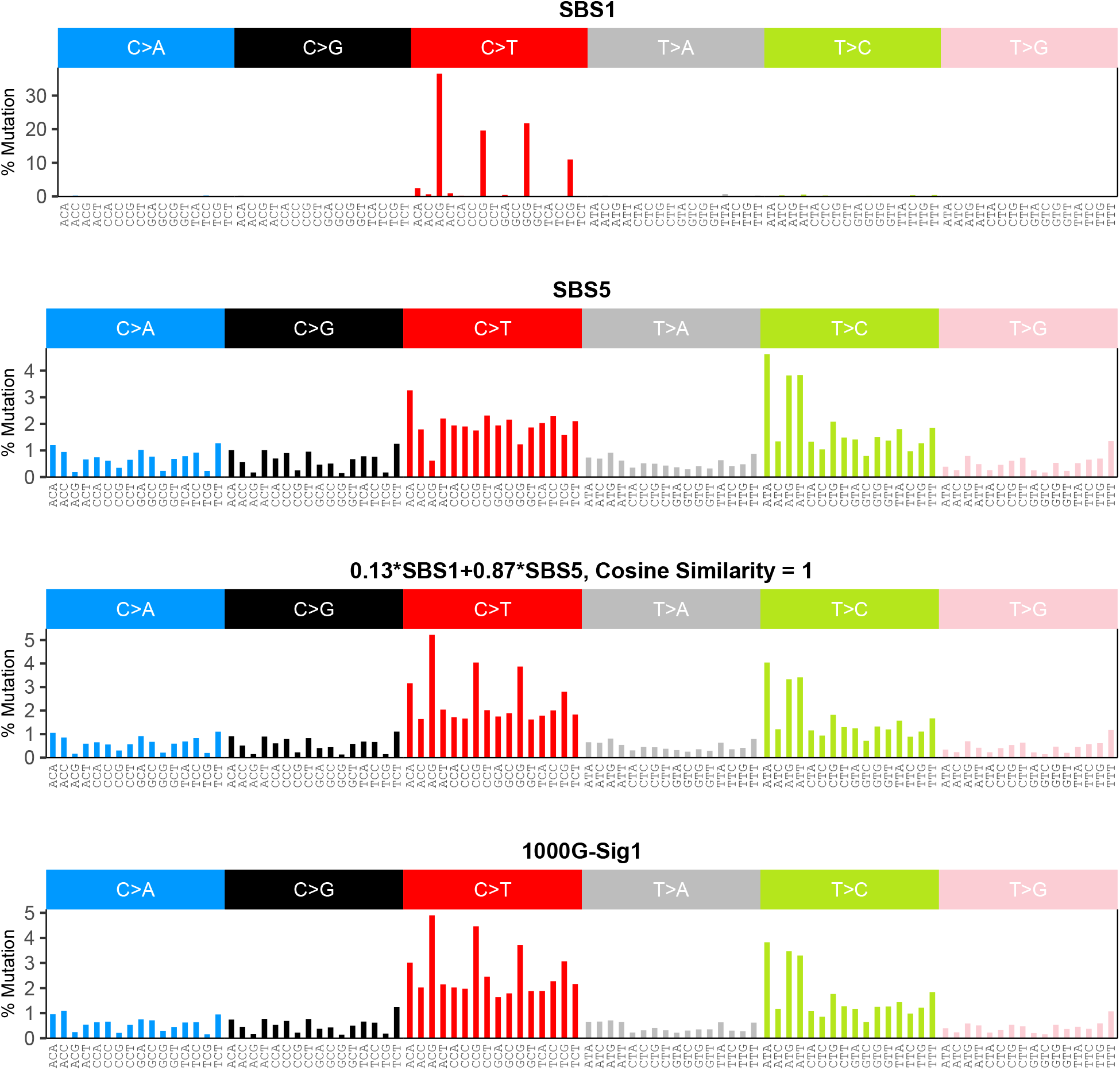
Reconstruction of 1000G-Sig1 using COSMIC signatures. The best model of 1000G-Sig1 is 0.13*SBS1+0.78*SBS5.

**Supplementary Figure S14.**
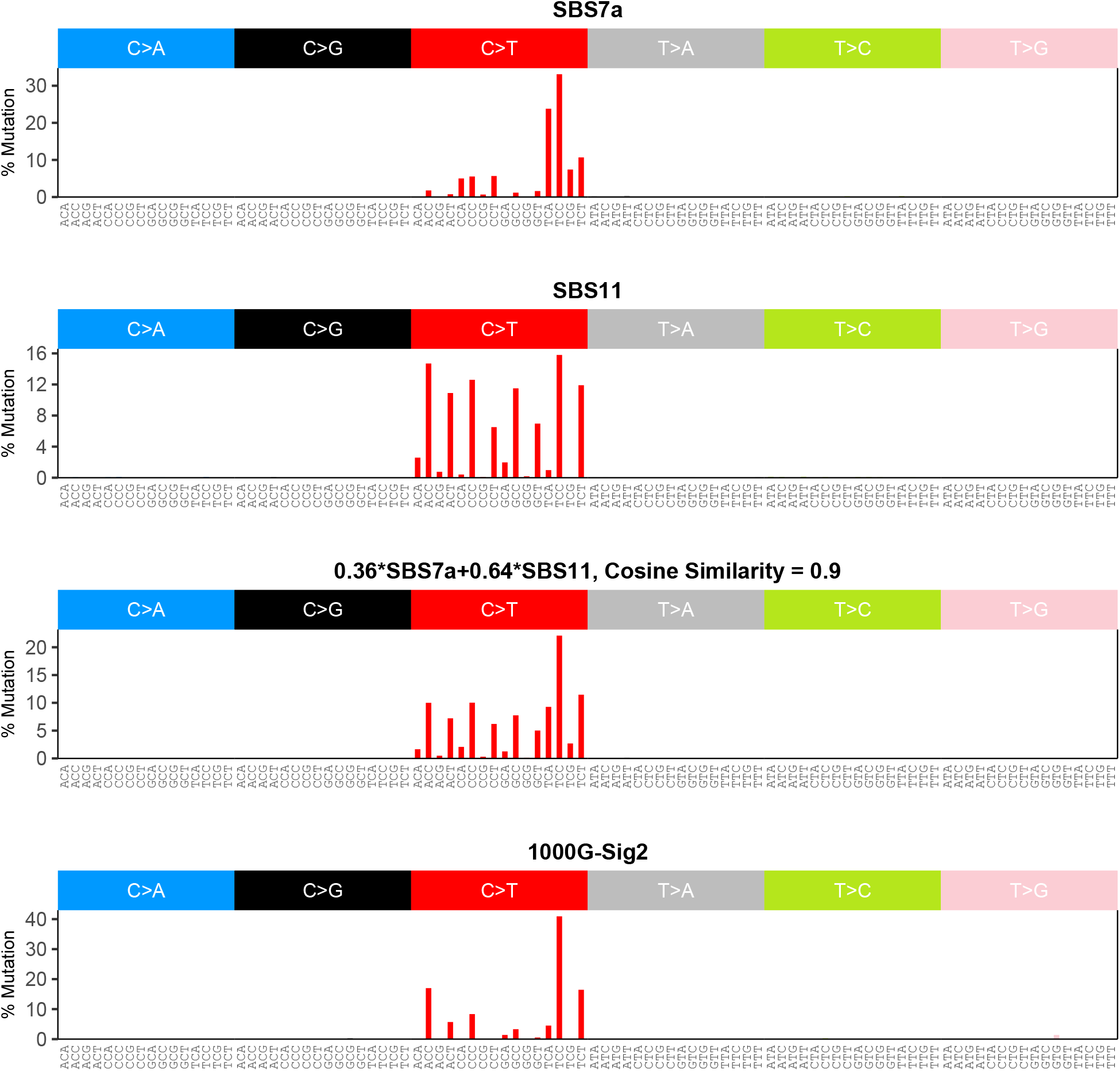
Reconstruction of 1000G-Sig2 using COSMIC signatures. The best model of 1000G-Sig2 is 0.36*SBS7a+0.64*SBS11.

**Supplementary Figure S15.**
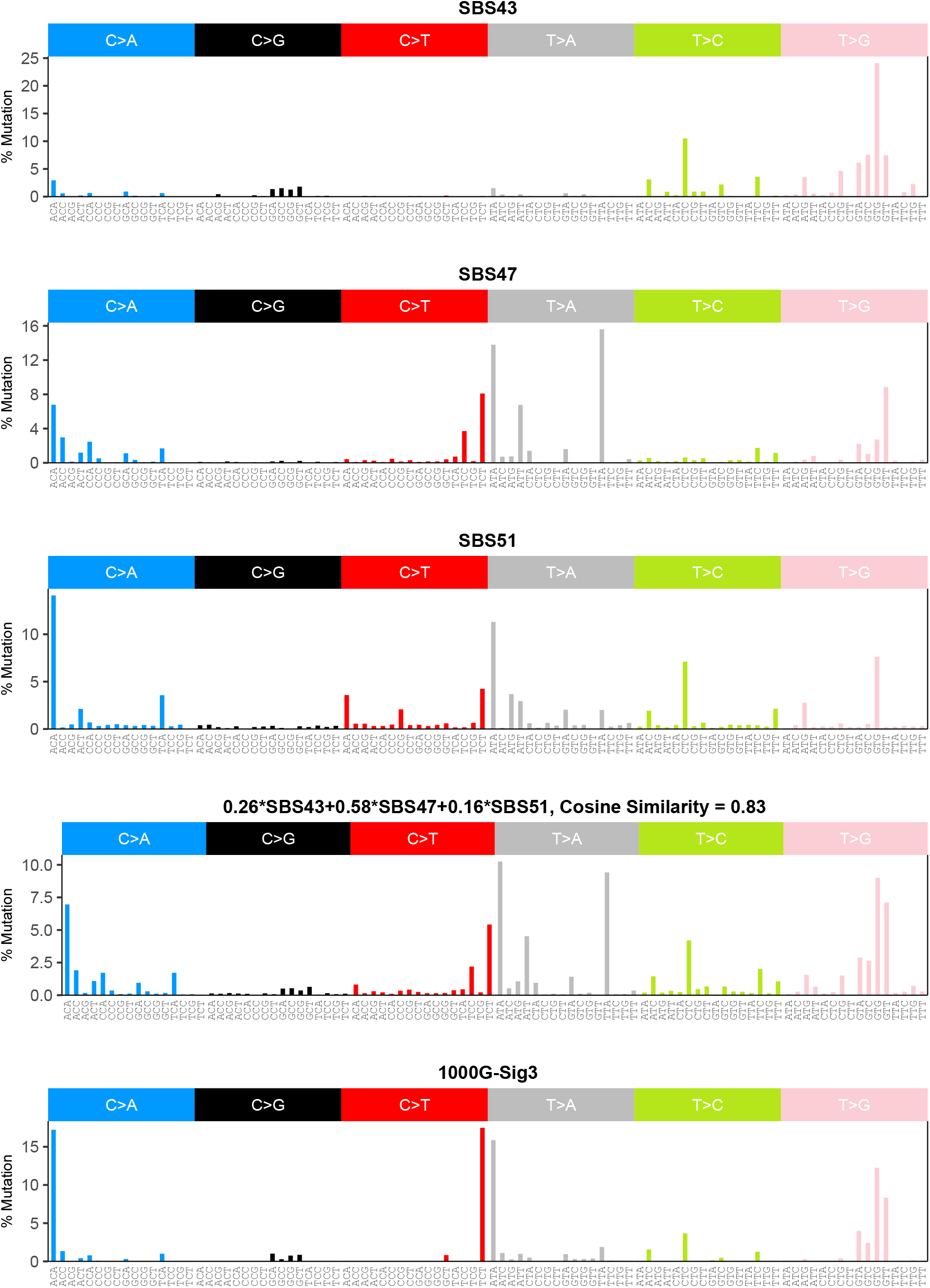
Reconstruction of 1000G-Sig3 using COSMIC signatures. The best model of 1000G-Sig3 is 0.26*SBS43+0.58*SBS47+0.16*SBS51.

**Supplementary Figure S16.**
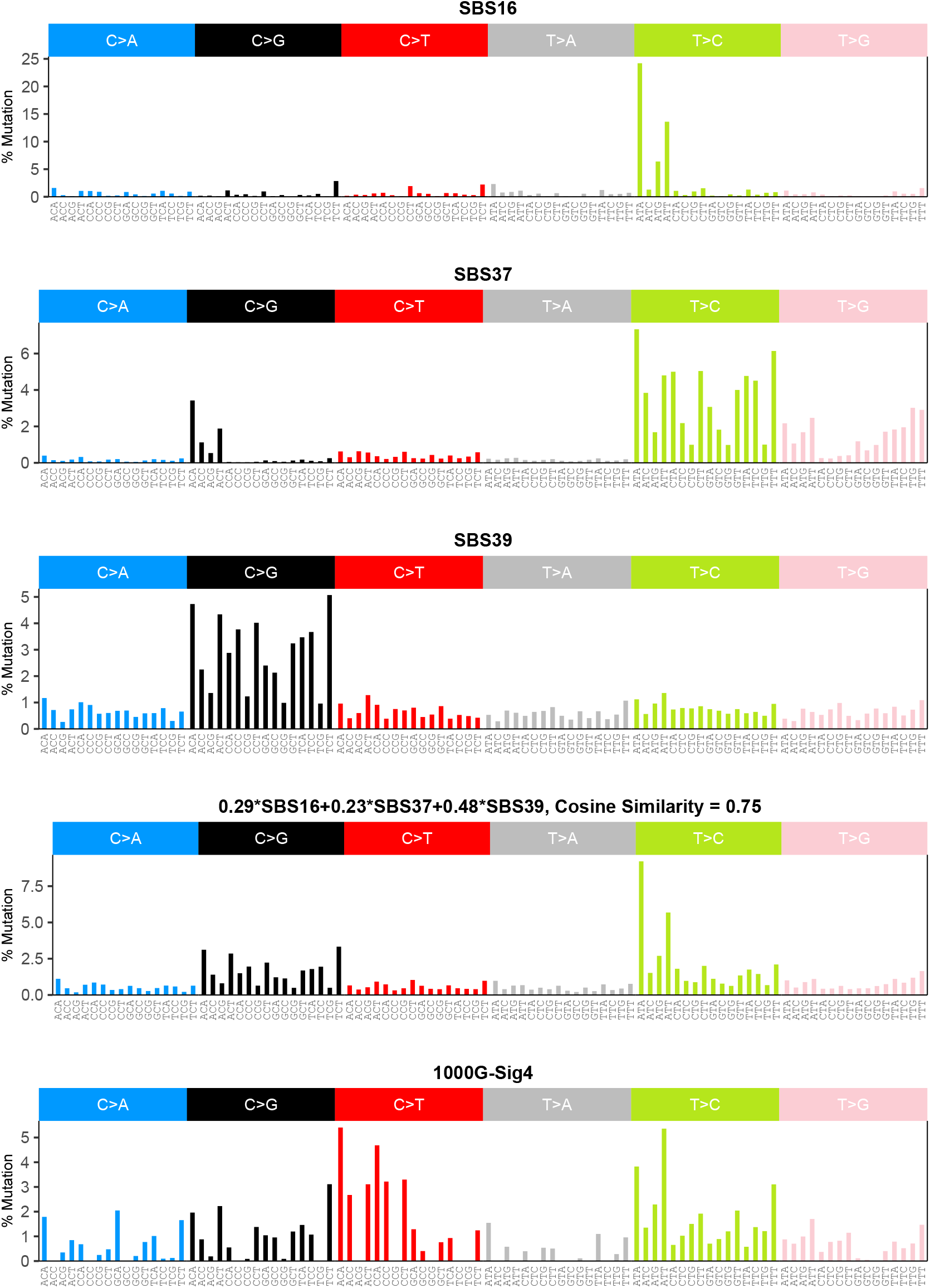
Reconstruction of 1000G-Sig4 using COSMIC signatures. The best model of 1000G-Sig4 is 0.29*SBS16+0.23*SBS37+0.48*SBS39.

**Supplementary Figure S17.**
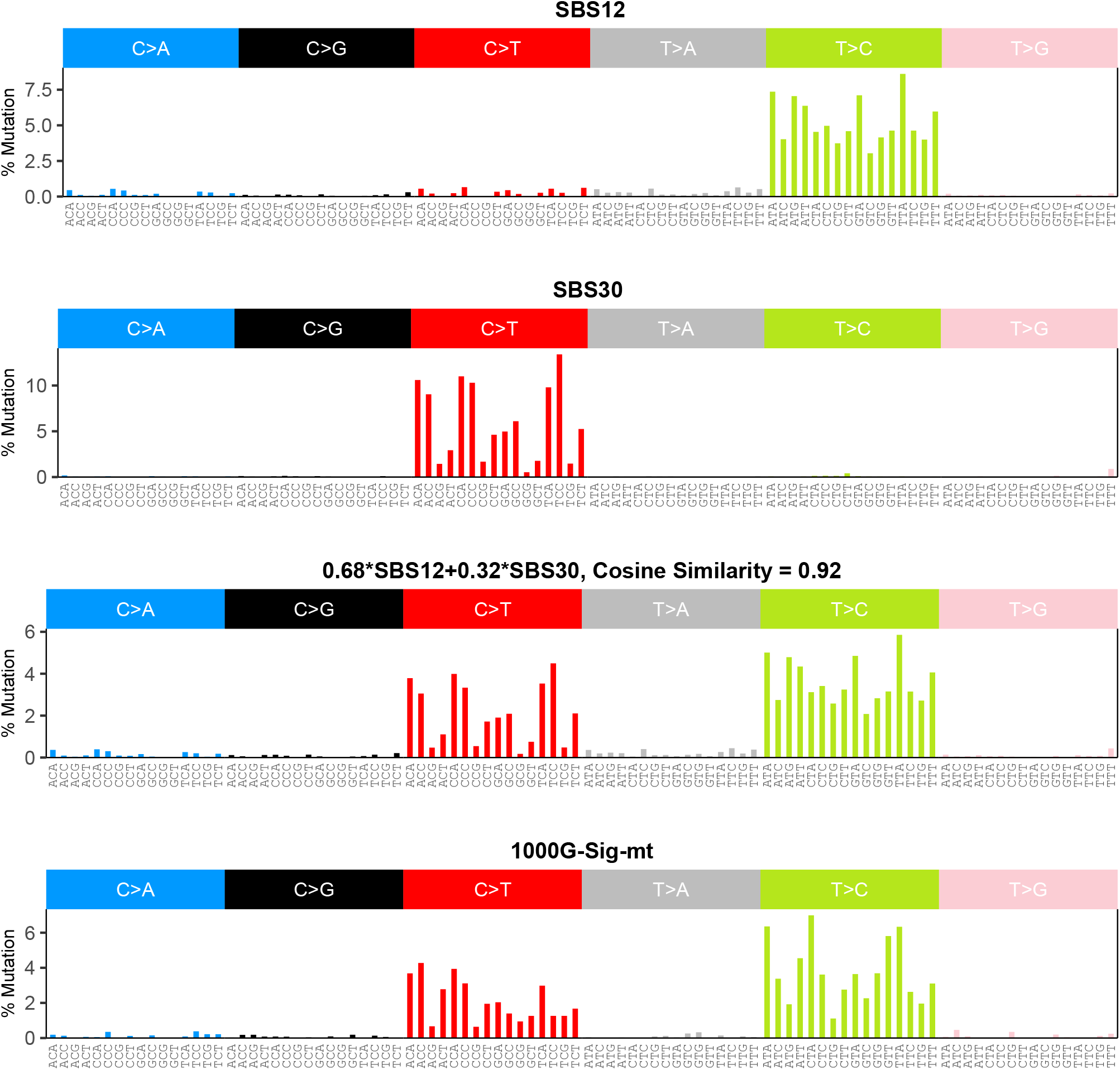
Reconstruction of 1000G-Sig-mt using COSMIC signatures. The best model of 1000G-Sig-mt is 0.68*SBS12+0.32*SBS30.

**Supplementary Figure 18.**
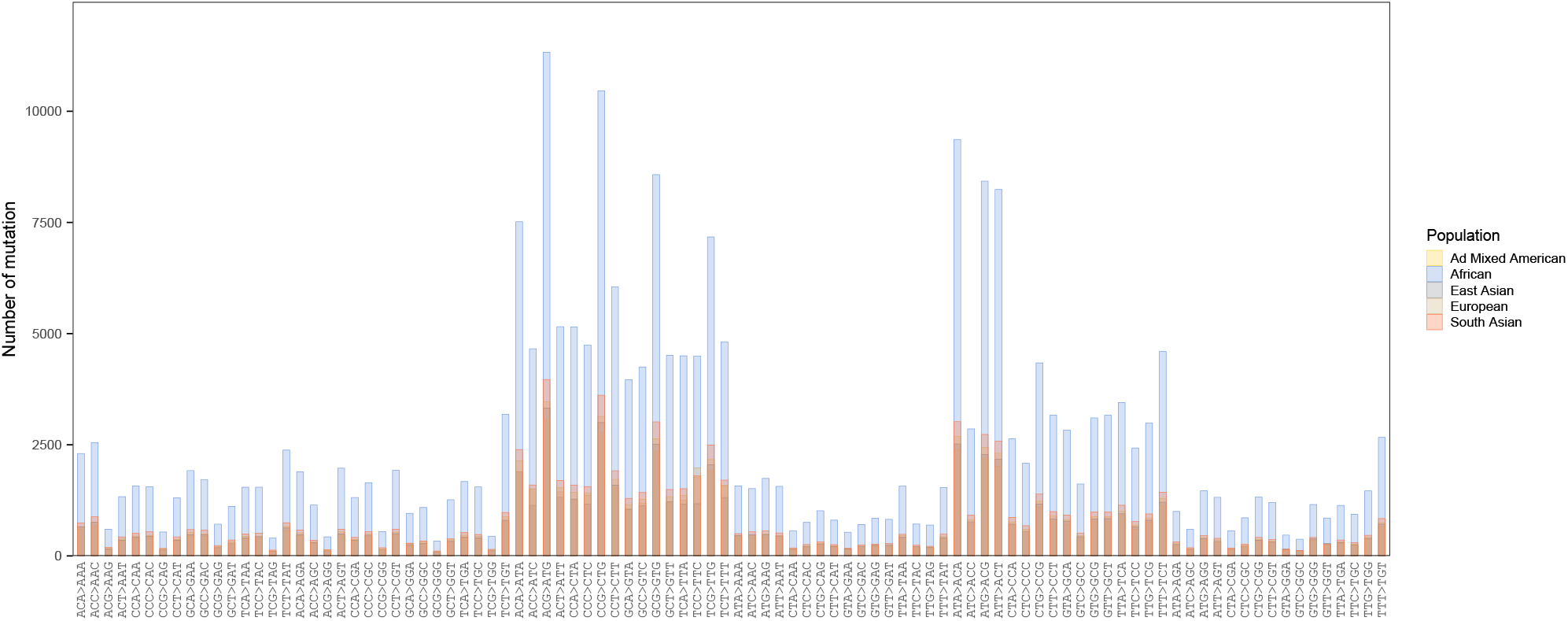
Analysis of 1000G Project datasets. Number of mutations (changes relative to the sequence of most common human ancestor) is shown for different human populations.

